# Brain regional identity and cell type specificity landscape of human cortical organoid models

**DOI:** 10.1101/2022.02.17.480833

**Authors:** M. Magni, B. Bossi, P. Conforti, M. Galimberti, F. Dezi, T Lischetti, X. He, R.A. Barker, C. Zuccato, I. Espuny-Camacho, E. Cattaneo

## Abstract

*In vitro* models of corticogenesis using mouse and human pluripotent stem cells (PSC) have greatly improved our understanding of human brain development and disease. Among these, 3D cortical organoid systems are able to recapitulate some aspects of *in vivo* cytoarchitecture of the developing cortex.

Here, we tested three cortical organoid protocols for brain regional identity, cell type-specificity and neuronal maturation. Overall all protocols gave rise to organoids that displayed a time-dependent expression of neuronal maturation genes such as those involved in the establishment of synapses and neuronal function. We showed that three months old cortical organoids showed a pattern of gene expression that resembled late human embryonic cortex. Comparatively, directed differentiation methods without WNT activation gave rise to the highest degree of cortical regional identity in brain organoids. Whereas, default “intrinsic” brain organoid differentiation produced the broadest range of cell types such as neurons, astrocytes and hematopoietic-lineage derived microglia cells of the brain. These results suggest that cortical organoid models produce diverse outcomes in terms of brain regional identity and cell type specificity and emphasize the importance of selecting the correct model for the right application.

## INTRODUCTION

The study of the human cortex during development is critical for a better understanding of how this rostral brain region evolves, including its neuronal connectivity and function as well as how it is affected in a range of human disorders. The ability to isolate and grow mouse and human embryonic stem cells (ESC) and the discovery of directed and undirected ways to generate neural cells has allowed the generation of a range of human brain cellular models ^1–3^. An important milestone in this respect was the original description of the derivation of mouse cortical cells from mouse ESC in the absence of any morphogens. This protocol constituted the first proof-of-principle that non-directed methods for neural differentiation generate mostly cells of the rostral neural tube, such as the cortex ^4–6^. Similarly, human ESC were later differentiated following default conditions to generate cortical progenitors and neurons ^7,8^. These findings showed that the absence of caudalizing factors was sufficient to induce cortical identity *in vitro.* In an effort to increase the number of cortical cells derived from human embryonic (hESC), and human induced pluripotent stem cells (hiPS) different conditions such as the addition of retinoic acid to enhance neuronal maturation, or the use of dual SMAD inhibitors, to block TGFβ and BMP signaling pathways, have been used to enhance neuroectodermal fate ^9–11^. Subsequent addition of morphogens such as WNT, BMP, FGF8 or SHH were shown to pattern areal identity within the telencephalon ^4,7,12–15^ or to caudalize areal identity towards posterior midbrain and hindbrain fates ^16,17^. These *in vitro* models recapitulated important aspects of *in vivo* brain development such as temporal patterning and human specific patterns of neuronal maturation ^5,7,11^. Although these two-dimensional (2D) telencephalic models proved to be invaluable in studying important aspects of human brain evolution and diseases ^4,7,14,15,18–23^, other aspects such as long-term cell polarity as well as neuronal migration and integration into neural networks could not be as easily investigated in these reductionists *in vitro* systems. In one approach, the developmental potential of mouse and human ESC-derived cortical cells has been tested after transplantion into the more suitable environment of the mouse brain *in vivo* ^5,7,15,24–27^. In another approach, *in vitro* tridimensional (3D) models of corticogenesis have been developed from human PSC that recapitulate to some extent *in vivo* cortical cytoarchitecture and cell polarity in a dish. Such models have provided a cellular environment that supports the formation of ventricle-like structures around which progenitors lie within a ventricular-like region (VZ) separated from a rudimentary cortical plate-like area ^28–32^. 3D organoid models are either based on default differentiation, where human neuroectodermal cells acquire a rostral forebrain identity in the absence of caudalizing signals ^28,33,34^, or based on the use of morphogenes that direct the differentiation towards specific neuronal subtypes ^29–31^. Guided-differentiation protocols incorporate a combination of different morphogens such as the use of dual SMAD inhibition to enhance neuroectodermal fate ^9,29,31^. This first ‘neural induction’ step is followed by an amplification step or proliferative step in the presence of EGF and FGF ^29^ or in the presence of a TGFβ inhibitor together with a GSK3β inhibitor, that results in activation of WNT signaling ^31^. Cortical organoid protocols may also use morphogens to enhance the maturation profiles; embed organoids in a matrix basement membrane-like structure to support polarity and cell growth; and/or use bioreactors to increase oxygen and nutrient diffusion into the tissue structures at late stages in the differentiation ^29–31,35^. However, to date it remains unclear which conditions are necessary and sufficient to generate specific brain cell types, to best recapitulate *in vivo* cell polarity and to generate mature neuronal populations.

Here, we explored the differentiation capacity of three cortical 3D organoid models, one of them based on default differentiation and the use of a matrix basement ^28^ and two guided differentiation protocols with or without matrix inclusion of the organoids in the absence or presence of WNT activation ^29,31^. We analysed the organoids at the early progenitor and late neuronal stages and identified their brain regional identity based on the expression of regional-specific genes from the rostro-caudal and dorsal-ventral axis of the neural tube. Furthermore, we screened the organoids for the expression of neuronal maturation genes and compared them with human embryonic and adult cortical brain samples. Finally, we compared the various models for their ability to generate different brain cell types such as oligodendrocytes, astrocytes and non-neuroectodermal derivatives such as microglia cells.

## RESULTS

### Intrinsic and directed-differentiation cortical organoid models show a high efficiency of neuronal specification

We differentiated H9 cells using either a default “intrinsic protocol” in the absence of any morphogenes ^28^(PC1), a directed differentiation protocol following dual SMAD inhibition and GSK3β inhibition/WNT activation coupled to TGFβ inhibition ^31^ (PC2) and a directed differentiation protocol with dual SMAD inhibition followed by EGF and FGF treatment ^29^ (PC3) (Figure 1A). H9 cells were aggregated and exposed to default or directed differentiation followed by either a (i) matrigel embedded step from 2 weeks onwards (PC1), (ii) a matrigel embedded step for about 1 week (PC2) or (iii) no matrigel-embedding step (PC3) (Figure 1A). These protocols were adapted to the use of an orbital shaker from 2 weeks *in vitro* until the last time point in culture to promote diffusion of oxygen and nutrients inside the organoids, as an alternative to the use of a spinning bioreactor (see materials and methods) ^36^. After the neuroectodermal induction step, organoids were transferred to neuronal medium and left to mature until 3 months in culture without (PC1), or in the presence of neuronal maturation morphogens (PC2 and PC3) to accelerate and improve the outcome of mature neuronal populations (Figure 1A). We analysed organoids from PC1, PC2 and PC3 protocols at early neuronal progenitor (days 7, 25) and late neuronal stages (days 50, 90). As expected, the size of the organoids increased upon time (Figure S1A,E).

**Figure 1.**
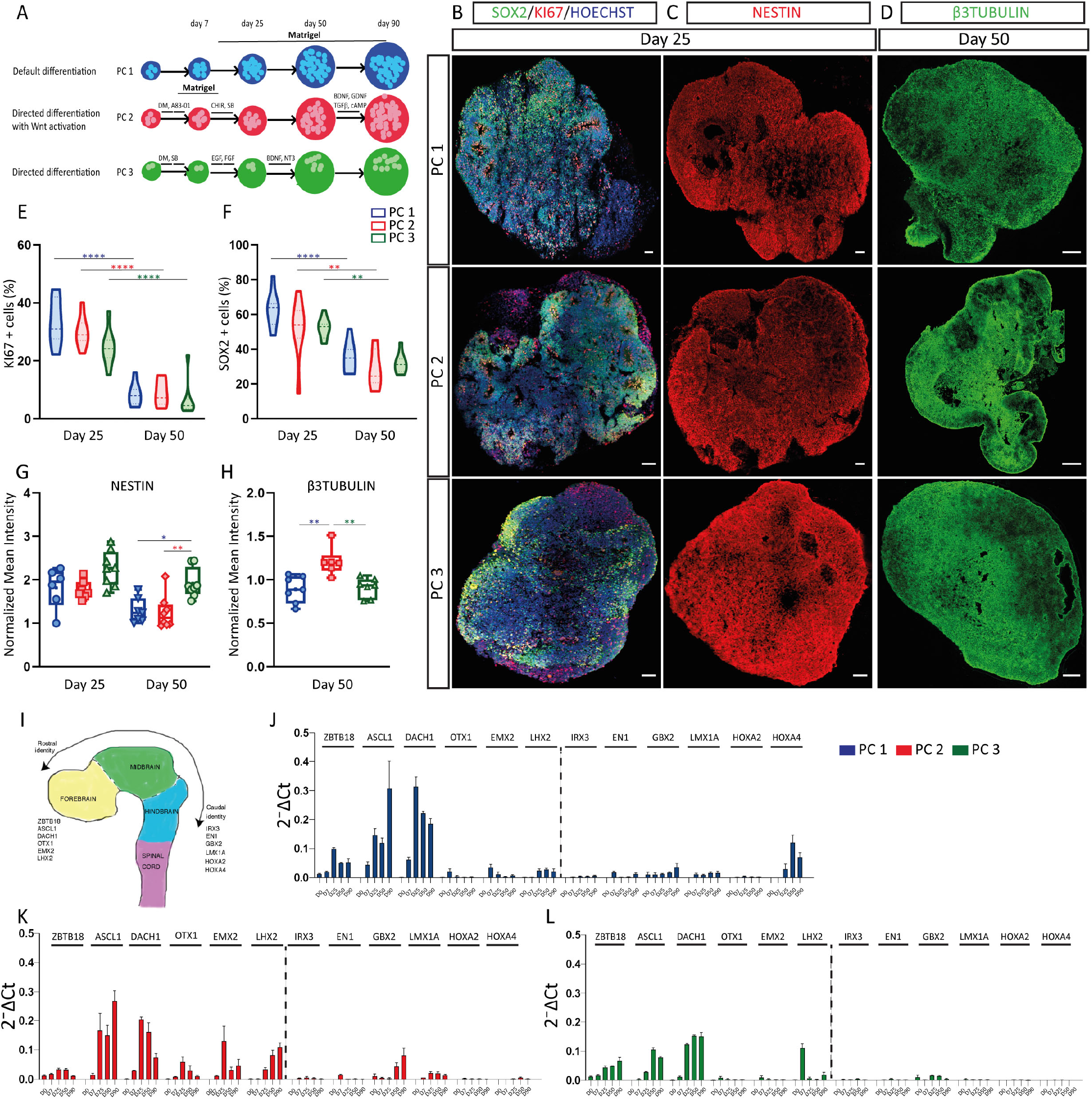
Default and directed differentiation cortical protocols efficiently generate neuronal progenitors and neurons. (A) Schematic representation of the experimental outlay. PC1 (default differentiation); PC2 (directed differentiation with WNT activation); PC3 (directed differentiation without WNT activation). Organoids were collected at early progenitor stages d7, d25 and late neuron stages d50, d90. (B-D) Immunofluorescence images showing the expression of the neuronal progenitor markers SOX2 (green; B); NESTIN (green; C); the proliferation marker KI67 (red; B); and the neuronal marker beta 3 tubulin (green; D) in day 25 and 50 organoids from PC1, PC2 and PC3 protocols. Scale bars represent 100 μm (B-D). Violin plots showing the percentage of cells expressing KI67 (E) and SOX2 (F), and box graphs showing the normalized mean intensity of NESTIN (G) and β3TUBULIN (H) in PC1, PC2 and PC3 organoids. n=8 sections from 4 organoids per protocol. Two way-ANOVA (E-G) and One-way ANOVA (H) with Tukey post-tests. *p < 0.05, **p < 0.01, ***p < 0.001, ****p < 0.0001. (I) Cartoon depicting the Rostral-Caudal axis of the developing neural tube. (J-L) Biomark analysis of PC1, PC2, PC3 organoids (days 7, 25, 50, 90) for the expression of rostral: ZBTB18, ASCL1, DACH1, OTX1, EMX2, LHX2, and caudal: IRX3, EN1, GBX2, LMX1A, HOXA2, HOXA4z neural tube genes. Data are shown as absolute normalized amount of mRNA (2-^ΔCt^) ± SEM, PC1 (n=3-9); PC2 (n=2-7); PC3 (n=3-4).

In order to better understand the differences between organoids derived from the three protocols we performed a high throughput quantitative PCR analysis by Biomark HD assay (Fluidigm) (see samples used in Tables S1 and S2 and raw data in Table S3). Organoids derived from the PC1 method showed the highest variability based on principal component analysis (PCA) (Fig. S1D), suggestive of a high variability in cell fate identity by default methods, as reported ^28,35,37^.

Stem cell renewal genes were, as expected, downregulated following differentiation. Interestingly, mesoderm germ layer markers such as CDX2 and KRT14 were enriched in PC1 and PC2-derived organoids, indicating the presence of a pool of non-neuronal cell types in these conditions ^38^ (Fig. S1F-H). The three protocols generated organoids that expressed neuronal progenitor and proliferative markers such as NESTIN, KI67 and SOX2. The percentage of neuronal progenitor and proliferative cells was similar between protocols and decreased from early progenitor stage (day 25) to later time points (day 50) (Fig. 1B-C, E-G and Fig. S1B). We detected broad expression of the radial glia marker Blbp and the neuronal markers doublecortin (DCX) and β3Tubulin (β3TUBB) in organoids from all protocols (Fig. 1D,H, Fig. S1H and Fig. S2).

Overall these data indicate that the three protocols generate organoids with overall expression of neuronal progenitor and neuronal markers, suggesting an efficient neuroectodermal acquisition and neurogenesis in all conditions.

### Directed differentiation without WNT agonists generates organoids with the highest cortical identity

Next, we analysed brain regional identity among cortical organoid protocols by comparing the expression of brain rostro/caudal and dorso/ventral-specific genes (Fig. 1I and Fig. S3A). We observed enrichment in rostral regional markers in organoids from all three protocols. In contrast, caudal brain regional markers were less abundant but more represented in PC1 and PC2 organoids (Fig. 1J-L). Analysis of dorsal and ventral telencephalic identity revealed an overall enrichment of dorsal genes and expression of some lateral ganglionic eminence (LGE)-specific genes *versus* medial ganglionic eminence (MGE) identity genes (Fig. 2 and Fig. S3). PAX6, the early neuroectodermal fate determinant and dorsal progenitor marker was detected from early stages in all three protocols with the highest expression in PC2-derived organoids (Fig. 2 and Fig. S1C), suggestive of higher neuroectodermal induction following dual SMAD inhibition, as reported ^9^. Strikingly, we found a higher proportion of TBR1+ neurons for deep layers of the cortex in PC3-derived organoids and higher proportion of CTIP2+ (deep layers) and SATB2+ (upper layers) neurons in PC1 and PC3-derived organoids (Fig. 3A-G). Telencephalic and cortical-specific markers such as FOXG1, EMX1, TBR2, NEUROD1, TBR1 were more expressed in PC3-derived organoids at three months (day 90), consistent with a higher degree of cortical identity of PC3-derived organoids (Fig. 3H). Moreover, representation of the top 10 up Biomark genes from each protocol using the human Brain Atlas database showed a comparative enrichment of human cerebral cortex and basal ganglia-specific genes for PC3 organoids, and a broader and more caudal regional identity for PC1 and PC2 conditions (Fig. 3I-K).

**Figure 2.**
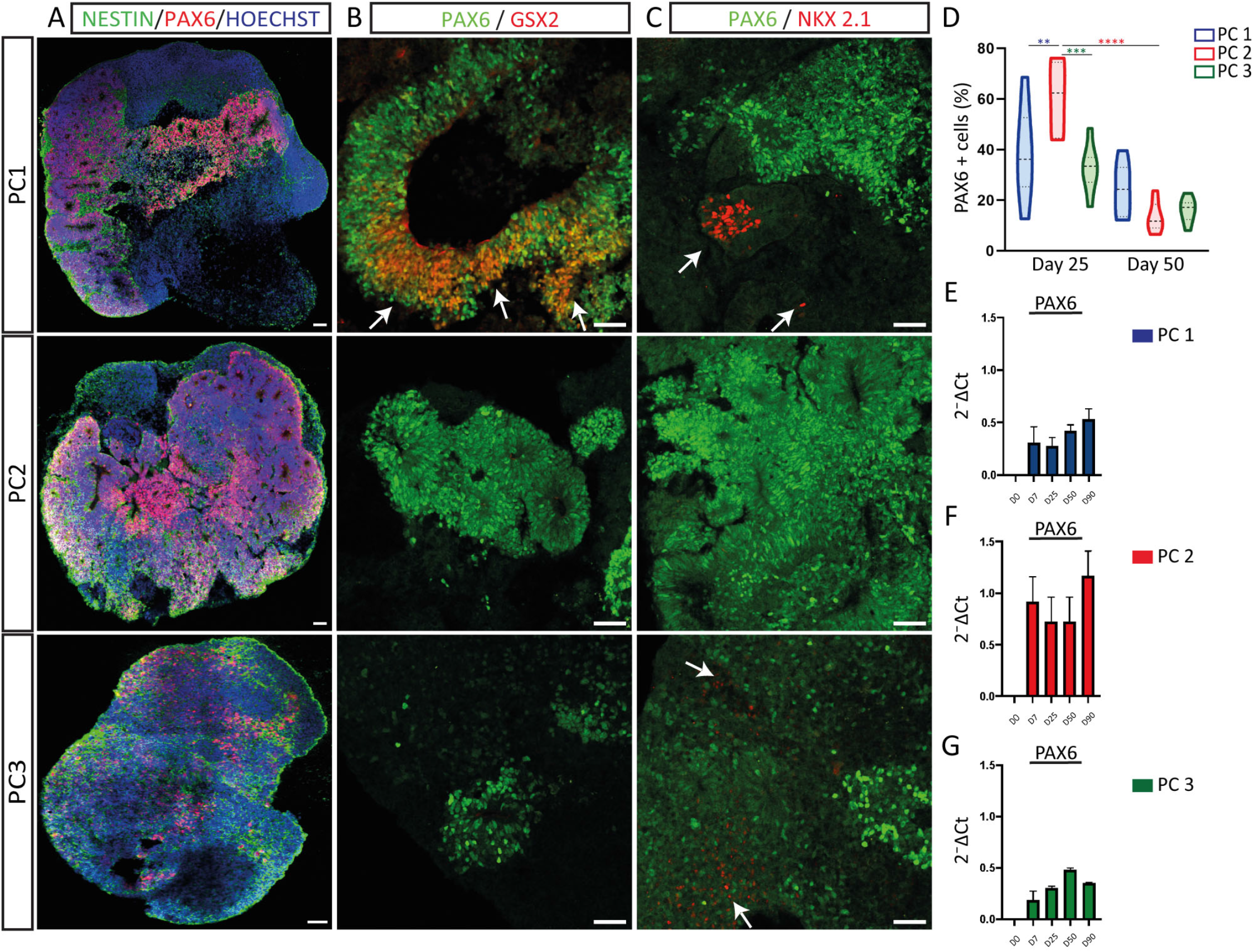
Cortical organoid differentiation protocols give rise to dorsal and ventral telencephalic identity progenitor populations. (A-C) Immunofluorescence images showing the expression of the neuronal progenitor marker NESTIN (green; A), the dorsal progenitor marker PAX6 (A-C), the ventral LGE and MGE progenitor marker GSX2 (red; B) and the MGE progenitor marker NKX2.1 (red; C) in day 25 organoids from PC1, PC2 and PC3 protocols. Counterstaining of nuclei was done with Hoechst (blue). Arrows show the presence of ventral telencephalic GSX2 (B) or NKX2.1 (C) positive progenitor cells in PC1 and PC3 organoids. Scale bars represent 50 μm. (D) Violin plot showing the percentage of cells expressing Pax6. n=8 sections from 4 organoids per protocol. Two way-ANOVA with Tukey post-tests. **p < 0.01, ***p < 0.001, ****p < 0.0001. (E-G) Biomark analysis of PC1, PC2, PC3 organoids (days 7, 25, 50, 90), for the expression of the dorsal forebrain-specific gene PAX6. Data are shown as absolute normalized amount of mRNA (2-^ΔCt^) ± SEM, PC1 (n=3-9); PC2 (n=2-7); PC3 (n=3-4).

**Figure 3.**
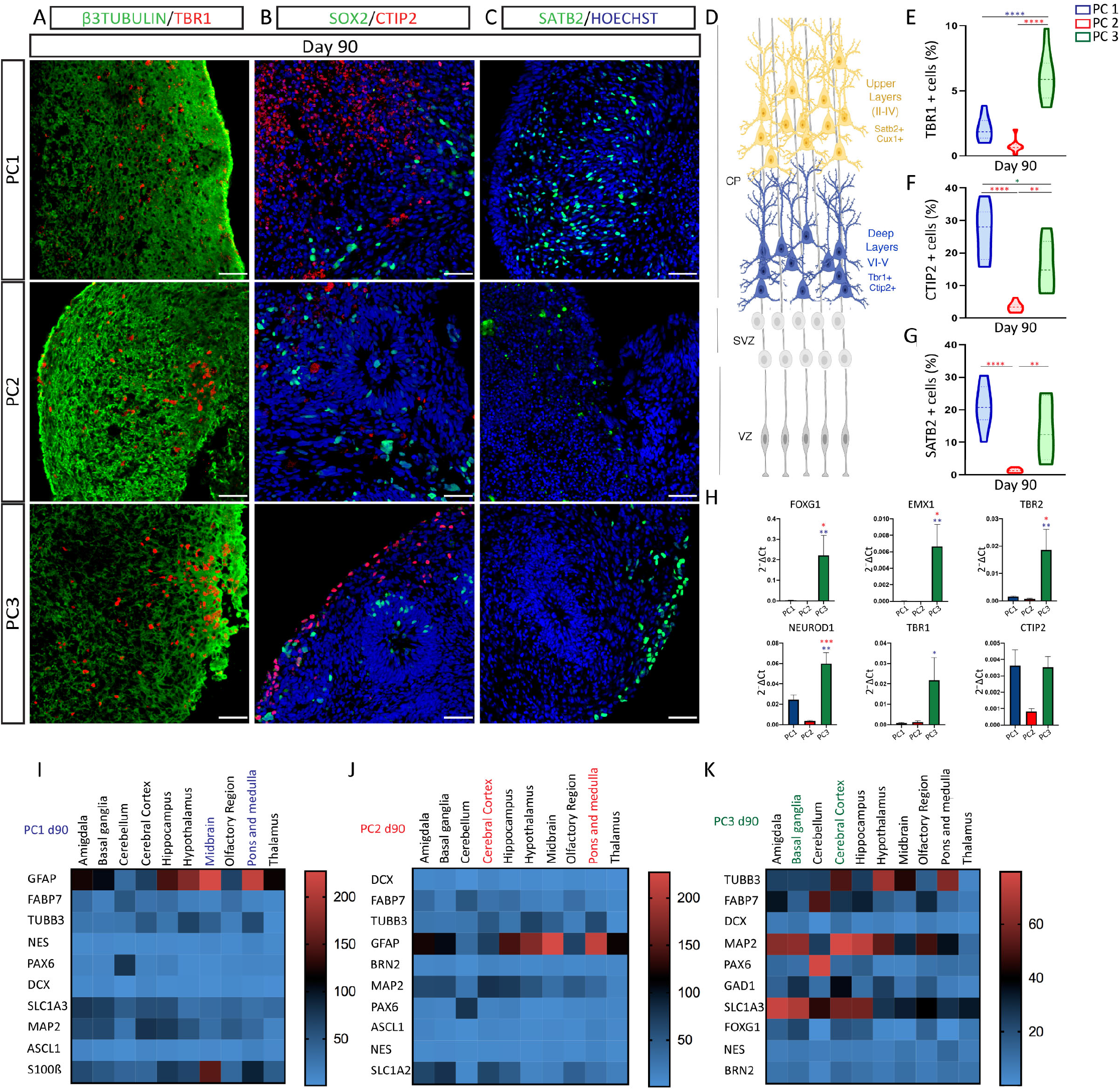
Directed differentiation without WNT agonists generates the highest cortical-specific progenies. (A-C) Immunofluorescence images showing the expression of the neuronal marker β3TUBULIN (green; A); the neuronal progenitor marker SOX2 (green; B); the deep layer neuronal markers TBR1 (red; A) and CTIP2 (red; B); and the upper layer neuronal marker SATB2 (green; C) in organoids following PC1, PC2 and PC3 protocols. Counterstaining of nuclei was done with Hoechst (blue). Scale bars represent 50 μm. (D) Cartoon depicting the different progenitor regions (VZ, SVZ) and neuronal compartments (CP: deep layers; upper layers) of the developing cortex. (E-G) Violin plots showing the percentage of cells expressing TBR1 (E); CTIP2 (F) and SATB2 (G) in PC1, PC2 and PC3 organoids. n=8 sections from 4 organoids per protocol. One way-ANOVA with Tukey post-tests. *p < 0.05, **p < 0.01, ****p < 0.0001. (H) Biomark analysis of PC1, PC2, PC3 organoids (days 7, 25, 50, 90), for the expression of cortical-specific genes: FOXG1, EMX1, TBR2, NEUROD1, TBR1, CTIP2. Data are shown as absolute normalized amount of mRNA (2-^ΔCt^) ± SEM, PC1 (n=3-9); PC2 (n=2-7); PC3 (n=3-4). One way-ANOVA with Tukey post-tests. *p < 0.05, **p < 0.01, ***p < 0.001. (I-K) Heatmap representation of the expression of the top 10 up Biomark genes from PC1 (I), PC2 (J) and PC3 (K) organoids in human brain samples (amigdala, basal ganglia, cerebellum, cerebral cortex, hippocampus, hypothalamus, midbrain, olfactory region, pons and medulla, thalamus) from Brain Atlas database: https://www.proteinatlas.org/humanproteome/brain.

These data suggest that all protocols generate organoids containing rostral brain regional populations, where directed differentiation without WNT activation shows the highest enrichment of cortical identity progenies.

### 3D organoid differentiation protocols closely mimic *in vivo* cell polarity and cortical cytoarchitecture

*In vitro* organoid models of corticogenesis from hESC recapitulate the presence of polarized structures with an apical and basal side resembling the *in vivo* ventricular zone (VZ) of the developing cortex. In order to compare cell polarity among protocols we analysed day 25 organoids for the localization of progenitor markers in apical *vs* basal compartments and day 90 organoids for the separation of progenitor and neuronal markers. We first analyzed the localization of RG mitotic cells within VZ-like regions. We detected mitotic phospho-vimentin positive RG cells close to Pals1 + apical areas in organoids derived from the three protocols (white arrows; Fig. 4A) mimicking *in vivo* cell polarity in the VZ ^28,32^. Interestingly, the area of rosette-like neuroepithelia and VZ-like regions were smaller in PC3-derived organoids at day 25 (Fig. 4A-D,F-G). These differences suggest an effect of matrigel on the growth of neuroepithelia and cell polarity. Accordingly, radial orientation of RG Nestin positive scaffold appeared most organized in PC1 when compared to PC2 and PC3 organoids (Fig. 4B, inset). Interestingly, PC1-derived organoids showed the highest localization of KI67+ and SOX2+ progenitor cells inside VZ-like regions, suggestive of a better polarized structure following long-term matrigel-embedding (Fig. 4C-D,H-I) ^28,30,39^. CTIP2 positive cortical neurons were detected outside of the VZ-like region in day 90 organoids from all protocols (Fig. 4E), indicating a correct separation of progenitors and neurons, reminiscent of the *in vivo* compartmentalization of proliferative areas and cortical plate.

**Figure 4.**
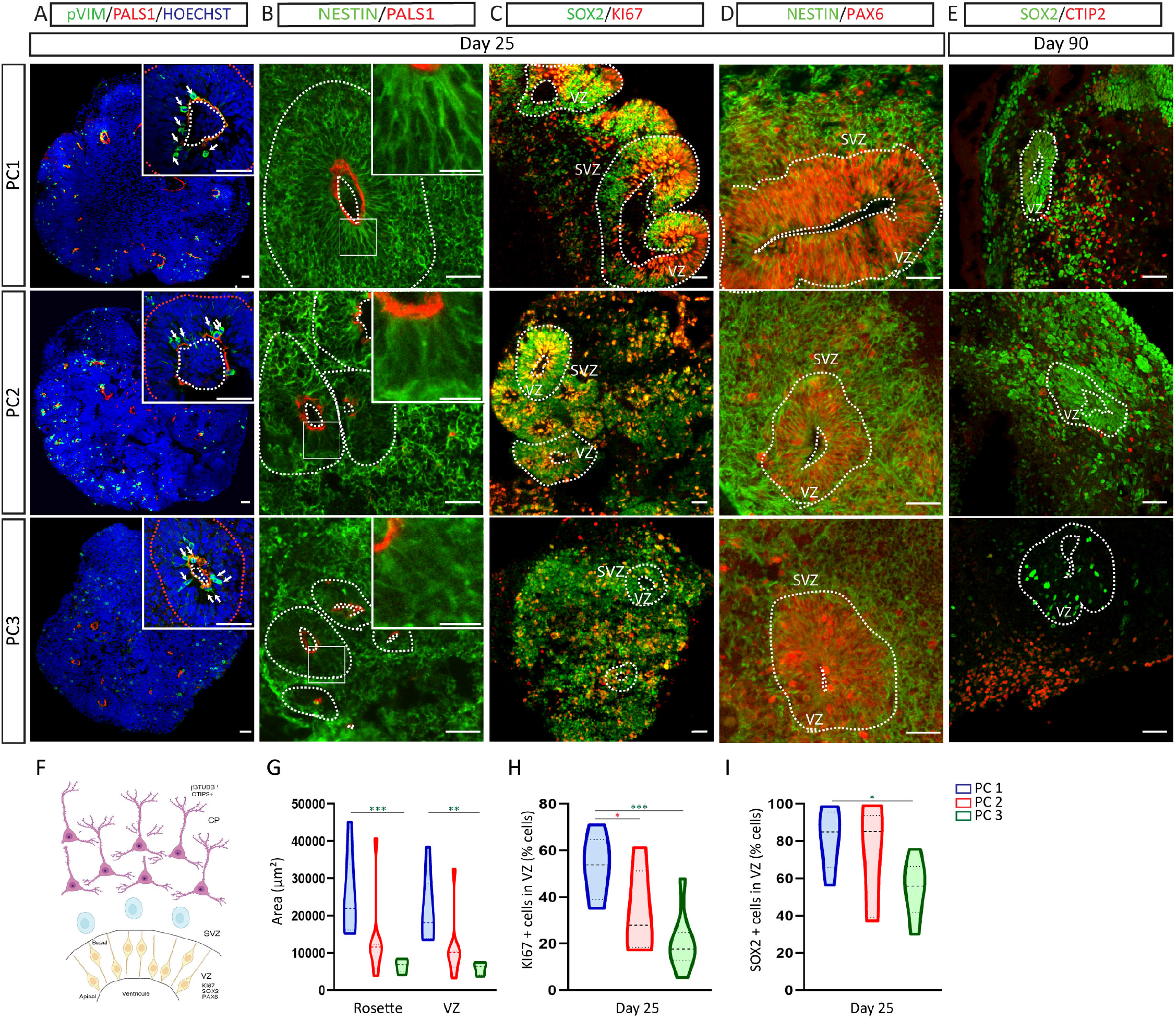
Protocols for cortical organoid differentiation recapitulate important aspects of cortical *in vivo* polarity. (A-D) Immunofluorescence images showing the expression of the apical marker PALS1 (red; A,B); the mitotic marker pVimentin (green; A); the neuronal progenitor markers NESTIN (green; B,D) and SOX2 (green; C); the proliferative marker KI67 (red; C) and the cortical progenitor marker PAX6 (red; D) at day 25. (E) Immunofluorescence images showing the expression of the neuronal progenitor marker SOX2 (green); and the deep layer neuronal marker CTIP2 (red) at day 50. Counterstaining of nuclei was done with Hoechst (blue). Dashed lines delineate the VZ-like proliferative regions in the organoids and the apical region of the VZ. Arrows show the presence of numerous pVimentin mitotic cells located at the apical side of the PALS1+ VZ (A). VZ, ventricular zone; SVZ, subventricular zone. Scale bars represent 100 μm (A), 50 μm (B-E), and top right insets 400 μm (A), 100 μm (B). (F) Cartoon depicting the apical-basal localization of radial glia cells within the VZ and the progenitor and neuronal cytoarchitecture of the developing telencephalon. (G) Violin plot showing the area (μm^2^) of neural rosettes and VZ-like areas. (H-I) Violin plots showing the percentage of cells expressing KI67 (H), and SOX2 (I) inside VZ-like regions from PC1, PC2 and PC3 organoids. n=8 sections from 4 organoids per protocol. One way-ANOVA with Tukey post-tests. *p < 0.05, **p < 0.01, ***p < 0.001.

These data show that all cortical organoid models mimic *in vivo* cell polarity and overall cytoarchitecture of the developing embryonic cortex where PC1 organoids display the most conserved cell polarity *in vitro*.

### Long-term culture of cortical organoids leads to the differentiation of mature neuronal populations

Human neurons derived from ESC and iPSC have been shown to follow a human species-specific program for maturation characterized by a very slow acquisition of mature neuronal cortical features whether in 2D *in vitro* paradigms or following transplantation into the developing mouse brain *in vivo* ^7,10,11,26^. In order to compare the degree of neuronal maturation among protocols we assessed the expression of several mature neuronal markers in organoids through time. We observed an increase in the number of NEUN+ and MAP2+ neurons and in the expression and number of VGLUT1+ puncta upon time in organoids derived from all protocols (Fig. 5A-F and Fig. S4A-C). The expression of mature neuronal genes, postsynaptic genes and glutamatergic genes were similarly upregulated in a time-dependent fashion (Fig. 5G-I). Interestingly, PC3 showed the highest level of expression of VGLUT1 among protocols, consistent with a higher degree of glutamatergic cortical identity (Fig. 5C,F-I and Fig. S4C).

**Figure 5.**
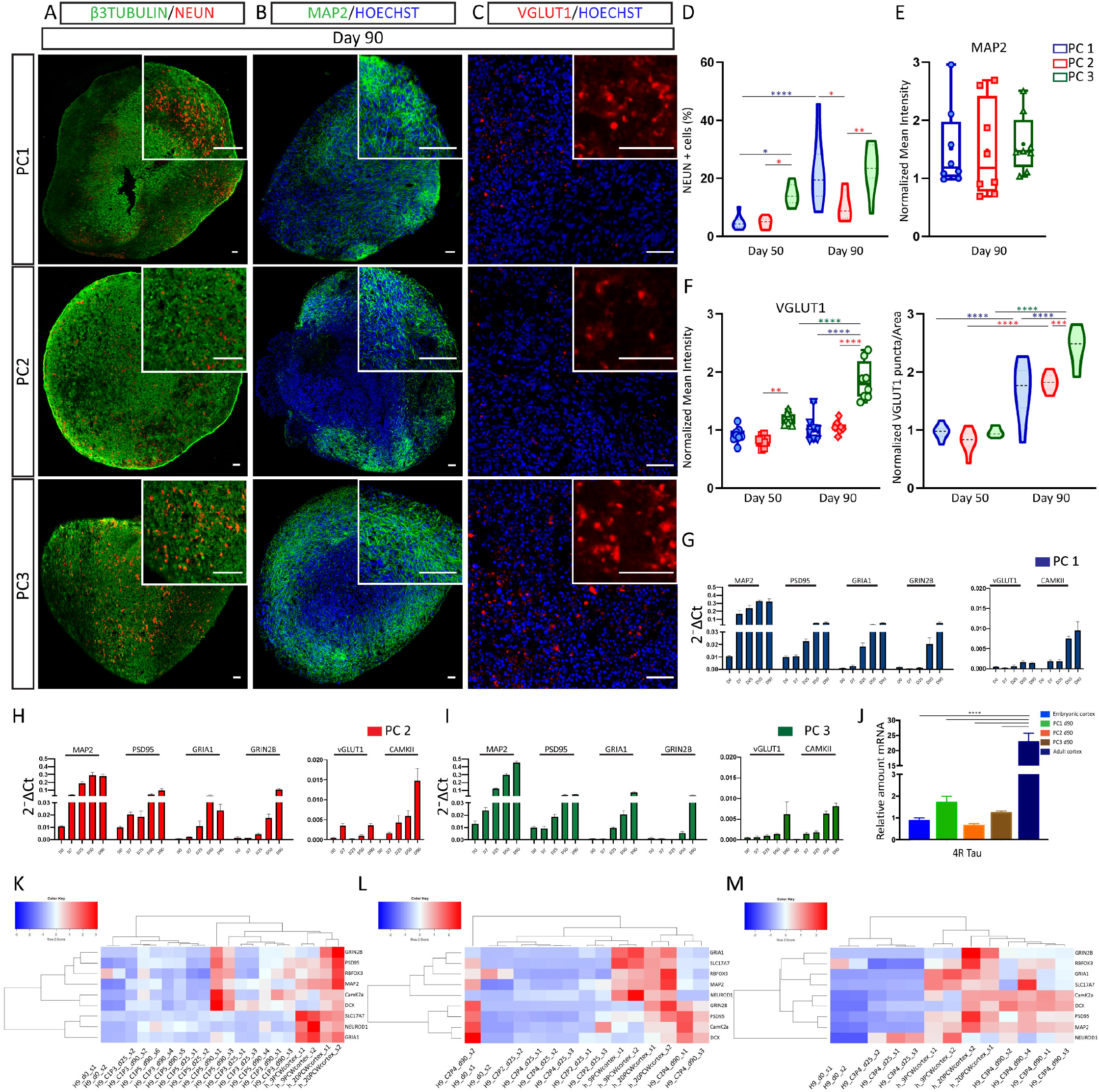
Long-term culture of 3D organoids leads to the generation of mature neuronal populations. (A-C) Immunofluorescence images showing the expression of the neuronal markers β3 TUBULIN (green; A); NEUN (red; A); MAP2 (green; B); VGLUT1 (red; C) in day 90 organoids from PC1, PC2 and PC3. Counterstaining of nuclei was done with Hoechst (blue). Scale bars represent 100 μm, and top right insets 400 μm. (D,F) Violin plots showing the percentage of cells expressing NEUN (D) and the normalized number of VGLUT1 puncta per area (F, right) in PC1, PC2, PC3 organoids. (E-F) Box graphs showing the normalized mean intensity of MAP2 (E) and VGLUT1 (F, left) in PC1, PC2, PC3 organoids. n=8 sections from 4 organoids per protocol. One way ANOVA (E) and Two way-ANOVA (D,F) with Tukey post-tests. *p < 0.05, **p < 0.01, ***p < 0.001, ****p < 0.0001. (G-I) Biomark analysis of PC1, PC2, PC3 organoids (days 7, 25, 50, 90) for the expression of neuronal maturation genes: MAP2, PSD95, GRIA1, GRIN2B, VGLUT1, CAMKII. Data are shown as absolute normalized amount of mRNA (2-^ΔCt^) ± SEM, PC1 (n=3-9); PC2 (n=2-7); PC3 (n=3-4). (J) qRT-PCR analysis of PC1, PC2, PC3 organoids (day 90), embryonic cortex and adult cortex for 4R TAU (MAPT) isoform expression. Data are shown as relative amount of mRNA compared to embryonic cortex, as value 1 ± SEM (fold change). PC1, PC2, PC3 n=3; embryonic cortex PCW9 n=2; and adult cortex (52-63 years) n=2. One way-ANOVA with Tukey post-test of all samples, ****p < 0.0001. (K-M) Cluster heatmap analysis showing the normalized expression of neuronal maturation genes in early (day 25) and late (day 90) PC1, PC2, PC3 organoids, two samples of undifferentiated H9 cells (day 0: s1,s2), and two samples from human embryonic cortex at 9 and 20 gestational weeks (9/20 GW) (s1,s2). Top: legend showing color code for comparative levels of expression.

Next, we compared the level of expression of the adult mature splice form of the MAPT gene (TAU), 4R TAU ^40^ in three months organoids, and human embryonic and adult brain samples. We confirmed that the pattern of expression of 4R TAU varies between embryonic and adult stage in humans (Fig. 5J) ^24,41^. The expression of 4R TAU in three months cortical organoids was similar to that of the human embryonic brain (Fig. 5J). Moreover cluster Heatmap representation of neuronal maturation genes revealed that late (day 90) brain organoid samples from PC1, PC2 and PC3 more closely resemble embryonic cortical samples (9PCW; 20PCW) than early (day 25) brain organoids or human ESC (day 0) (Figure 5K-M). Accordingly, the top 30 Biomark genes from three months organoids revealed fetal brain identity and protocol-specific categories (PC1: prefrontal cortex and spinal cord; PC2: prefrontal cortex and midbrain; PC3: prefrontal cortex and cerebral cortex) (Fig. S4D). These results suggest that brain organoids show a time-dependent maturation with increasing expression of genes essential for glutamatergic neuronal function like glutamatergic transporters and subunits of AMPA and NMDA receptors. Cortical organoids at three months of differentiation were similar to late human embryonic cortex where PC3 shows the highest expression of mature glutamatergic cortical genes.

### Default differentiation generates brain organoids with a broad range of glial cell-subtypes

Models for cortical organoid differentiation have been suggested to generate glia derivatives such as astrocytes, oligodendrocytes, non-excitatory neuronal identities such as ventral telencephalic GABAergic cells and few mesodermal hematopoietic lineage-derived cells such as microglia cells ^28^,^29,31,38,42^. However to date it is unclear to which extent these cell types are generated and enriched in different protocols. In order to characterize the presence of these cell types we analyzed organoids for the presence of cell-specific subtype markers. We found significantly higher expression of GABAergic-specific genes in three months PC3-derived organoids, suggesting an enrichment in ventral rostral telencephalic identity in the absence of WNT agonists, as expected ^16,17^ (Fig. 6A,D,G and Fig. S5A). Next we assessed the presence of astrocytes, oligodendrocytes and non-neural mesodermal-derived microglia cell identities. We found increased expression of astrocyte and microglia markers in PC1-derived organoids (Fig. 6B-C,E-F,H-I and Fig. S5B,D), suggesting a higher presence of these glia cell types following default differentiation conditions. In contrast, even though some oligodendrocyte markers were detected in late stage organoids we could not identify the presence of double positive OLIG2+ NKX2.2+ late oligodendrocyte progenitor cells (OPC), or of mature oligodendrocytes (MBP+) in any of the conditions tested (Fig. S5C and data not shown). Comparison of the top 30 Biomark genes from three months organoids revealed 45% of common genes among protocols, 7% of genes-specific for PC1 (astrocyte-specific and caudal SNC identity); 7% of genes-specific for PC2 (neuronal maturation and caudal identity); and 17% of genes-specific for PC3 (telencephalic and cortical-identity) (Fig. 6J).

**Figure 6.**
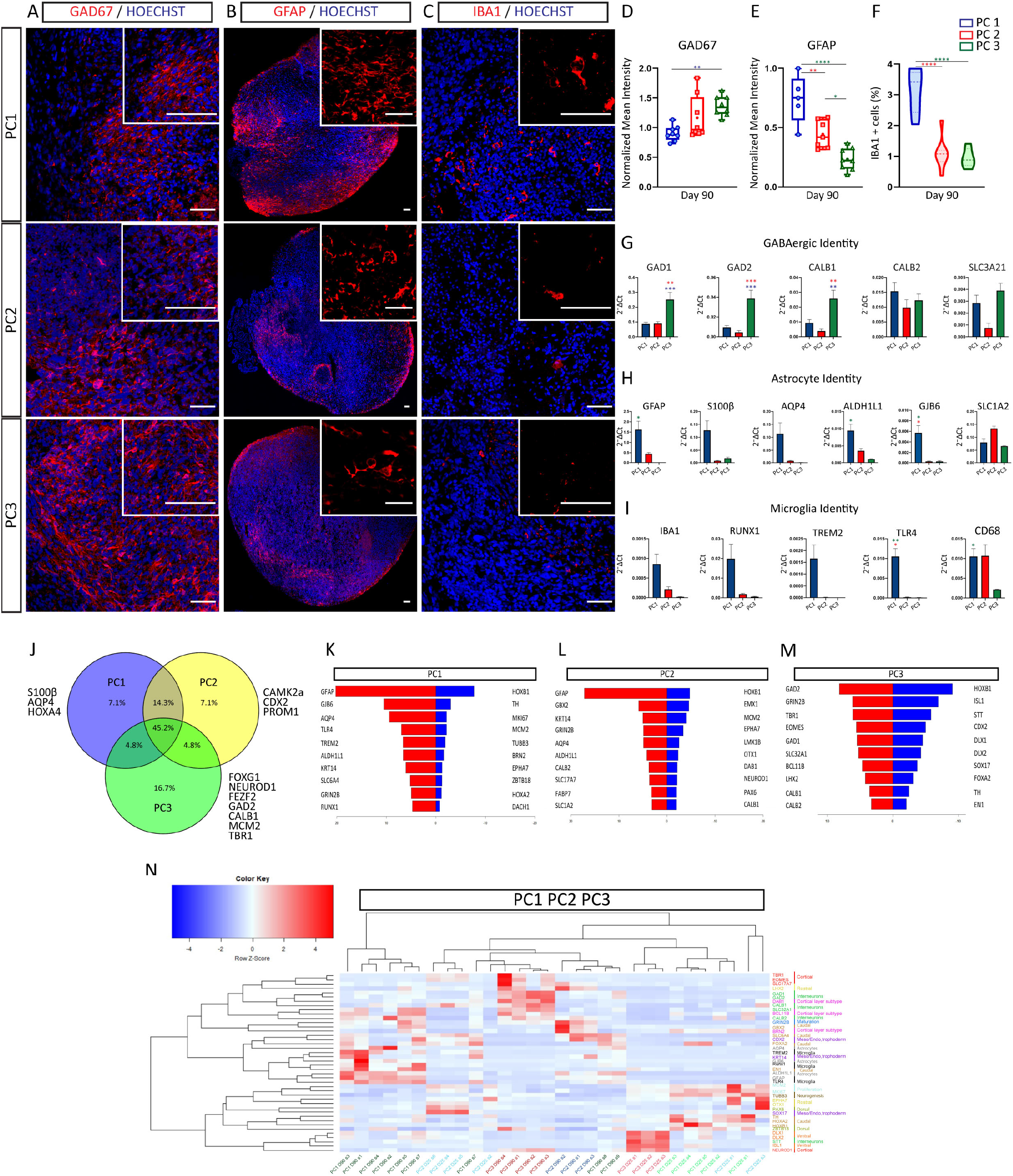
Cortical organoid differentiation protocols generate a diversity of neuronal and non-neuronal populations. (A-C) Immunofluorescence images showing the expression of the GABAergic neuronal marker GAD67 (red; A), the astrocytic marker GFAP (red; B) and the microglia marker IBA1 (red; C) in organoids following PC1, PC2 and PC3 protocols at day 90. Counterstaining of nuclei was done with Hoechst (blue). Scale bars represent 50 μm (A,C), 100 μm (B) and top right insets 100 μm (A,C) and 400 μm (B). (D-E) Box graphs showing the normalized mean intensity of GAD67 (D) and GFAP (E) in PC1, PC2, PC3 organoids. One way-ANOVA with Tukey post-tests. *p < 0.05, **p < 0.01, ****p < 0.0001 (F) Violin plot showing the percentage of cells expressing Iba1 in PC1, PC2, PC3 organoids. n=8 sections from 4 organoids per protocol. One way-ANOVA with Tukey post-tests. *p < 0.05, **p < 0.01, ****p < 0.0001. (G-I) Biomark analysis of PC1, PC2, PC3 organoids (days 7, 25, 50, 90) for the expression of GABAergic: GAD1, GAD2, CALB1, CALB2, SLC32A1; astrocytic: GFAP, S100β, AQP4, ALDH1L1, GJB6, SLC1A2; and microglia: IBA1, RUNX1, TREM2, TLR4, CD68 specific genes. Data are shown as absolute normalized amount of mRNA (2-^ΔCt^) ± SEM, PC1 (n=3-9); PC2 (n=2-7); PC3 (n=3-4). One way-ANOVA with Tukey post-tests. *p < 0.05, **p < 0.01, ***p < 0.001. (J) Venn diagram representing the top 30 up Biomark genes from PC1,PC2 and PC3 organoids. (K-M) Back-to-back representation of the biomark top 10 upregulated (red, left) and downregulated (blue, right) genes in late (three months) *vs* early (day 25) PC1, PC2 and PC3 organoids. (N) Heatmap analysis showing the normalized expression of the biomark top 10 upregulated genes from PC1, PC2 and PC3 in early (day 25) and late (day 90) organoids from all protocols. Color code genes: Cortical (red), Rostral identity (yellow), interneurons (light green), Cortical layer subtype (pink), neuronal maturation (dark blue), caudal identity (light brown), meso/endo, trophoectoderm (purple), astrocytes (grey), microglia (black), proliferation (light blue), neurogenesis (dark brown), dorsal (dark green) and ventral genes (orange).

Next, we analyzed the most differentially expressed genes between late (three months) and early brain organoids (day 25) to assess overall differentiation capacity among protocols. Interestingly, PC1 organoids top 10 upregulated genes at three months showed an upregulation of astrocyte, and microglia genes (Fig. 6K). PC2 organoids top 10 upregulated genes were in the category of astrocyte-specific as well as a caudal neural identity, whereas rostral and cortical specific genes were downregulated (Fig. 6L). We found that cortical-specific, ventral telencephalic and neuronal maturation genes were among the top 10 upregulated genes in three months PC3 organoids. Conversely, neuronal progenitor, caudal identity and proliferative genes were among the top 10 downregulated (Fig. 6M). Accordingly, when comparing the top 10 upregulated genes from each protocol across all organoids we observed that cortical-specific (red), cortical layer subtype (pink), rostral identity (yellow) and GABAergic identity (green) gene categories were best represented in PC3 late organoids (Figure 6N).

These data suggest that directed differentiation without WNT activation generates the highest enrichment of rostral telencephalic, both cortical glutamatergic and ventral GABAergic cell identities. Spontaneous differentiation on the other hand, leads to the highest number of glia cell-subtypes *in vitro*.

## DISCUSSION

Several 3D cortical organoid protocols have been published to date ^28–31,43^. Among those, default differentiation from human PSC has proven to be sufficient to generate cortical-like structures in the absence of any morphogens ^28,33,34^. Other protocols rely on an initial dual SMAD inhibition step to enhance neuroectodermal cell fate followed exposure to growth factors to promote neuronal maturation ^29–31,43^. In addition, some protocols follow embedding of human 3D brain structures into a basement membrane-like extracellular matrix only for one-week at an early stage ^31^; or from two weeks onwards ^28,44^, whereas others have grown organoids without scaffolds ^29,30,43^.

Here, we compared three methods for cortical organoid generation from human ESC based on directed or default differentiation strategies, with or without matrix scaffold, and we have analysed cell progeny identities in culture. We found that methods based on directed differentiation without WNT activation ^29,45^ generated the highest level of cortical cell identity. Furthermore, we found that addition of GSK3β inhibitors/ WNT signaling pathway activators in a directed differentiation protocol generates more caudal neural tube cell identities ^31^, as previously suggested in 2D models ^16,17^. Default differentiation conditions ^28,44^ generated organoids composed of the broadest range of brain areal identities and brain cell-subtypes, and showed the highest variability between organoids, as reported ^28,35,37^.

Conditions where cell polarity is best displayed and preserved at early and late time points in culture are essential factors to consider when choosing *in vitro* models suitable for the study of cell division, and neuronal migration defects in developmental diseases ^28,39,46^. To that end we compared the three protocols for the degree of polarity displayed in culture. We found a separation between progenitor regions and the localization of differentiated neurons in all organoid protocols. However, organoids embedded into matrigel generated the best organization of neuronal progenitors around ventricle-like structures with an extended radially oriented scaffold from radial glia cells. Furthermore, organoids embedded in matrigel scaffolds from about two weeks until the latest time point in culture resulted in the most elongated VZ-like neuroepithelium and polarized regions within organoids. Thus, these methods could be more suitable for the study of cell polarity of human cortical progenitors in normal development and in neurodevelopmental diseases.

An important feature for human cortical *in vitro* models is the degree of neuronal maturation reached in culture. Previous studies using 2D models have shown that human ESC-derived cortical neurons display a slow maturation program highly reminiscent of the human species. Indeed, 2-3 months old cultures have shown an expression profile comparable to that from human fetal brain at midgestational stages ^7,10,11^. Studies comparing 3D and 2D paradigms have suggested a slightly faster maturation profile of brain-like structures following organoid models ^29,47^. However, single cell RNAseq analysis revealed that the cellular composition of 3D organoids after 3, and even 6 months in culture still appears to be similar to the human fetal brain rather than postnatal or adult stages ^35,37^. More recently, protocols have used modified conditions such as the use of bioreactors or slicing the organoids, in an attempt to improve neuronal maturation and survival of brain structures after months *in vitro* ^30,31,34,35^. Despite all this, the associated *in vivo* age of organoids has remained similar to gestational week 19-24 of the developing human fetal brain ^29,31,34,35,37,48^. We compared the neuronal maturation profile across time in organoids derived from the three protocols. We observed that protocols for default differentiation or directed differentiation resulted in similar patterns of neuronal maturation profiles based on the expression of a variety of genes present in mature neuronal populations such as NEUN, and MAP2 structural gene; genes important for the formation and maintenance of synapses like PSD95; genes involved in synaptic plasticity like CAMKII; and genes involved in neuronal activity like GRIA1, GRIN2B and vGLUT1. As expected, the pattern of neuronal maturation was increased through time culminating at 3 months in culture. We looked into the expression of a specific splice form of the MAPT gene enriched in adult brain, the 4R TAU ^24^, which is associated with different tauopathies such as frontotemporal dementia and Alzheimer’s disease ^49^. We found a similar expression among organoids from different conditions, which matched that found in human embryonic, rather than adult, cortex. All of these findings support a late embryonic identity of the organoid samples after three months in culture. It would be interesting in future work to explore the conditions that could further accelerate aging in 3D organoids *in vitro* such as the use of factors that inhibit telomerases, the expression of truncated proteins from patients showing accelerated aging, such as progerin from Hutchinson-Gilford progeria syndrome (HGPS), the inhibition of autophagosome function, or the induction of DNA damage ^50–52^.

A general aim of 3D organoid models *versus* reductionist 2D paradigms is to generate a more physiological cellular environment that can closer recapitulate essential aspects of normal *in vivo* cortical development. Cortical development is a fine tuned process where different cellular types interact with each other and balance cell proliferation, cell cycle exit, neuronal migration and neuronal maturation ^53^. In this aspect it would be interesting to define the experimental conditions that more closely generate the broad spectrum of cell identities present in the cortex at different developmental stages. Most models of cortical organoid formation have shown the presence of additional brain cell types such as ventral telencephalic GABAergic interneurons, astrocytes, and some degree of oligodendrocytes and non-neuroectodermal derived cells such as hematopoietic-lineage derived microglia cells of the brain ^28,29,31,38,42^. However, due to the diversity of protocols and experimental conditions it is currently unclear which paradigms lead to a higher enrichment of these particular cell populations. We noticed absence of late oligodendrocyte precursor cells and mature oligodendrocytes in organoids derived from all protocols even at the latest time points tested. These results could indicate that longer periods in culture beyond three months are necessary to generate oligodendrocyte populations. This observation could reflect an intrinsic timing of oligodendrocyte generation in the dorsal telencephalon related to postnatal stages *in vivo* ^37^. An alternative explanation is that some factors essential for oligodendrocyte generation or maturation are lacking in the *in vitro* conditions. Of interest in this respect are recent studies using modified protocols for the generation and enrichment of oligodendroglia cell populations through exposure of neuroectodermal progenitors to EGF, FGF2, SHH agonists and oligodendroglia trophic factors that have been shown to be required for their maturation ^42,54^. We found however that directed differentiation in the absence of WNT agonists generated organoids with the highest number of GABAergic neurons, corresponding to a higher rostral forebrain identity. Interestingly, intrinsic default differentiation showed the highest yield of glial cell types such as astrocytes and hematopoietic-lineage derived microglia cells, in line with what was seen histologically ^28,35,37,38^. This suggests that default differentiation methods could be essential to recapitulate cell diversity in a dish to best study the brain in health and disease.

In summary we show that cortical organoid models generate a wide variety of cell identities corresponding to different brain areal identities and brain cell types that relates to the use of default or directed differentiation conditions. Moreover, we detected changes in cell polarity and/or size of VZ-like regions according to the presence or absence of extracellular matrix scaffolds. Our results also highlight that cortical organoids in culture display a time-dependent neuronal maturation that nevertheless remain similar to that from the embryonic cortex, rather than to adult cortical samples even after three months. Our findings point out the importance of selecting the right cortical organoid differentiation method according to the areal identity, brain cell-subtype and cell polarity degree required for each specific brain model.

## Supporting information

Supplementary data Magni et al 2022

## ACKNOWLEDGEMENTS

We thank Massimiliano Pagani and Marco de Simone for the use and help with the Fluidigm-Biomark HD system. We thank Vittoria Bocchi from the University of Milan, and Aranaud Lavergne from the Bioinformatics team of GIGA-Genomics from the University of Liege, Belgium for their help and advice with the data analysis.

## CONFLICT OF INTEREST

The authors declare no competing interests.

## AUTHOR CONTRIBUTIONS

I.E.-C., M.M, B.B, performed all of the differentiation experiments and analysis with the help of M.G.. P.C. performed the Biomark assay. F.D. and T.L. performed the bioinformatics analysis of data. H.X., R.A.B. and C.Z. provided the human embryonic and adult brain samples. I.E.-C. wrote the manuscript with the help of all coauthors. I.E.-C. and E.C. designed and analysed all of the experiments.

## ETHICS APPROVAL

All procedures related to human tissues were approved by the research ethical committees and research services division of the University of Cambridge and Addenbrooke’s Hospital in Cambridge in accordance with the Human Tissue Act 2006 and by the Ethics Committee of the University of Milano, and ethics approval was obtained on 27 March 2013.

Tissue was handled in accordance with ethical guidelines and regulations for the research use of human brain tissue set forth by the National Institute of Health (NIH) (http://bioethics.od.nih.gov/humantissue.html) and the World Medical Association Declaration of Helsinki (http://www.wma.net/en/30publications/10policies/b3/index.html).

## FUNDING

This work was funded by the European Commission H2020 projects Neuromics Grant 305121 and NeurostemcellRepair Grant 602278 (to E. Cattaneo), and by the European Program Horizon 2020 Marie Sklodowska-Curie Actions MSCA-I.F.-2017 (to I. Espuny-Camacho).

## METHODS AND MATERIALS

### Human ESC Differentiation into 3D cortical organoid models

Cortical organoids were generated with few modifications (see supplemental materials and methods) from previous published methods (PC1: ^28^; PC2: ^31^; PC3: ^29^).

### Quantification of percentage of cells, mean intensity and number of puncta per area

Stained organoid sections were imaged by confocal microscopy (TSC SP5, Leica Microsystems) with a 20x objective to image Z series stacks of areas inside human organoids. All images were acquired using identical acquisition parameters as 16-bit, 1024×1024 arrays. Percentage of cells, mean intensity and number of puncta per area were quantified using Fiji/ImageJ software.

Percentage of cells (%) and number of puncta was calculated using a default threshold and generating masks to identify particles. Number of puncta per area was calculated by dividing the number of identified puncta among the area. Mean intensity of IF images were subtracted to background values. Number of puncta per area and mean intensity values were related to an internal control value of 1. Data are represented as Mean values. Statistical analysis was done with two way-ANOVA with Tukey post-tests.

### Quantification of organoid size

Organoid size was analysed at days 25 and 90 *in vitro* organoids. Fixed, cryopreserved and cryosectioned organoids (12-15 μm sections) were imaged following nuclear staining using Hoechst using a Leica DMI6000B microscope (Leica Microsystems) with an air 4x objective driven by Las-AF software. Image J/Fiji software was used to evaluate the area of the organoid pictures and represented as mm^2^.

Days 25 and 90 PC1 (n=4 organoids); PC2 (n=4 organoids); PC3 (n=4 organoids). Statistical analysis was performed using a two-way ANOVA with Tukey post-tests, ns= non-significant.

### Human embryonic and adult cortical samples

Post-mortem human fetal cortical specimens from 9pcw were obtained from University of Cambridge, UK. All procedures were approved by the research ethical committees and research services division of the University of Cambridge and Addenbrooke’s Hospital in Cambridge in accordance with the Human Tissue Act 2006. RNA from post-mortem human brain specimens at 20pcw was obtained according to an established protocol ^55^. Post-mortem adult cortical samples were obtained from the Harvard Brain Tissue Resource Center (HBTRC) (Belmont, Massachusetts, USA). Tissue was handled in accordance with ethical guidelines and regulations for the research use of human brain tissue set forth by the National Institute of Health (NIH) (http://bioethics.od.nih.gov/humantissue.html) and the World Medical Association Declaration of Helsinki (http://www.wma.net/en/30publications/10policies/b3/index.html).

### Biomark HD assay (Fluidigm)

For high-content qPCR experiments, cDNA was pre-amplified using a 0.2X pool of primers prepared from the same gene expression assays as were used for the Biomark analysis. Gene expression experiments were performed using the 96×96 qPCR Dynamic Array microfluidic chips (Fluidigm). Housekeeping genes used for normalization of the data were YWHAZ and GADPH. Data were analyzed and gene expression was carried out through the use of ΔCT as normalized absolute gene expression analysis: 2^-ΔCt^ whereby ΔCt=Ct_target_-Ct_normalizer_. Statistical analyses were done using a One-way ANOVA test (Analysis Of Variance), with Tukey post-tests and indicated as * for p<0.05; ** for p<0.01; *** for p< 0.001; **** for p< 0.0001.

### Statistical analysis

Data from experiments was processed with GraphPad Prism v.7 software. Statistical analysis was done following a one-way ANOVA or two-way ANOVA test (Analysis Of Variance), with Tukey post-tests and indicated as * for p<0.05; ** for p<0.01; *** for p< 0.001; **** for p< 0.0001.

## Notes

### Competing Interest Statement

The authors have declared no competing interest.

