## Supplementary data Magni et al 2022 for "Brain regional identity and cell type specificity landscape of human cortical organoid models"

### Supplemental Materials

#### Supplemental Figures

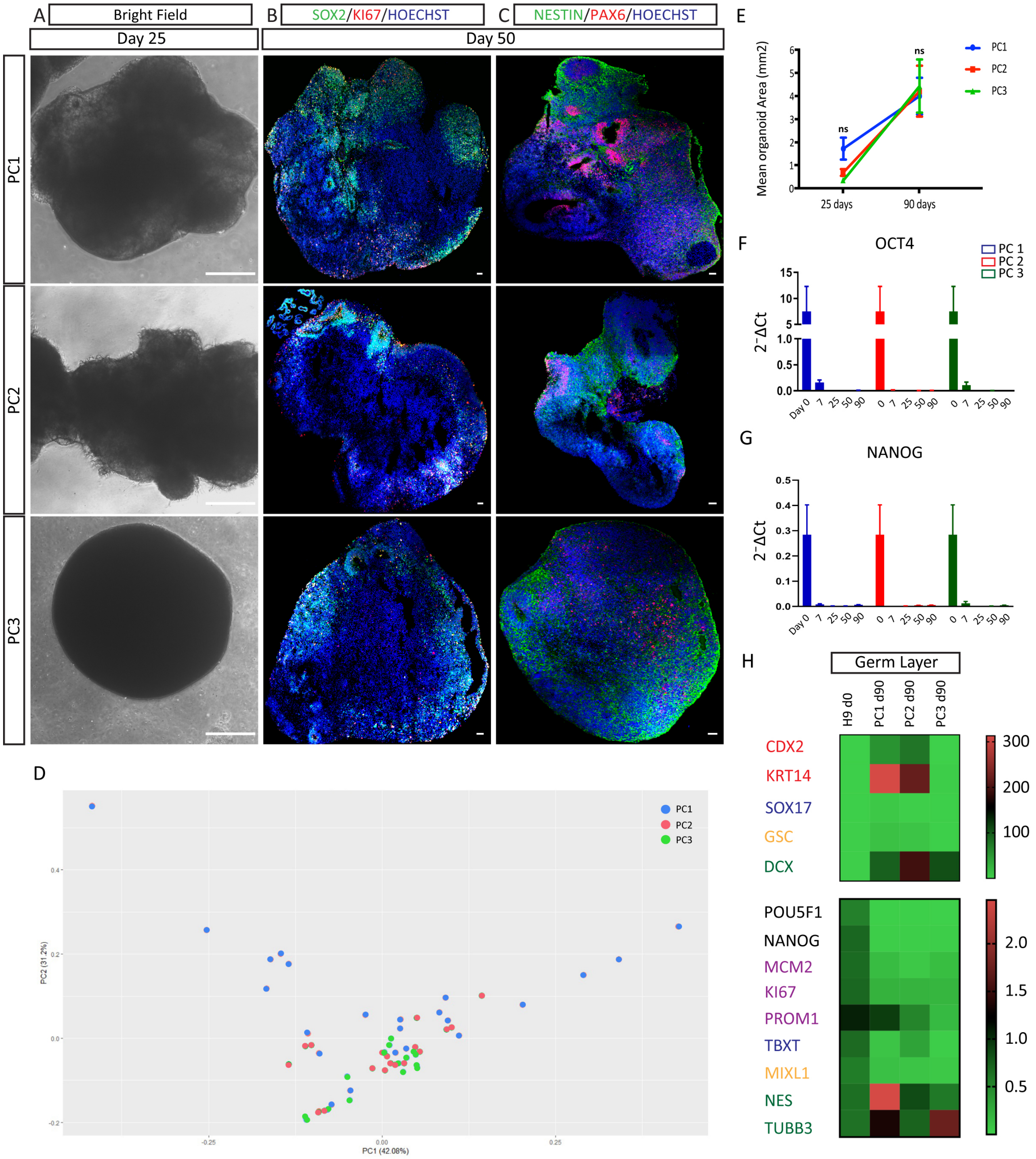

**Figure S1: Cortical organoids derived from intrinsic or directed-differentiation are composed of neuronal progenitors**

(A) Bright-field images of 3D organoids from PC1 (default differentiation), PC2 (directed differentiation with Wnt activation) and PC3 (directed differentiation w/o Wnt activation) at day 10-12 *in vitro*. (B-C) Immunofluorescence images showing the expression of the neuronal progenitor markers Sox2 (green; B); Nestin (green; C); the proliferation marker Ki67 (red; B); and the dorsal progenitor marker Pax6 (red; C) in day 50 organoids from PC1, PC2 and PC3 protocols. Scale bars represent 250 μm (A) and 100 μm (B-C). (D) Principal component analysis of all organoid samples (day 7, 25, 50, 90) from biomark analysis of 90 genes (PC1 in red; PC2 in green; PC3 in blue). (E) Quantification of organoid area (mm2) after 25 days and 90 days from wide-field images of cryosectioned organoids from PC1, PC2 and PC3. Data are represented as Mean ± SEM, PC1 (n=4); PC2 (n=4); PC3 (n=4), two-way ANOVA with Tukey post-tests, ns=non-significant. (F-G) Biomark analysis of PC1, PC2, PC3 organoids (days 7, 25, 50, 90) for stem cell genes Oct4 (E) an Nanog (F). Data are shown as absolute normalized amount of mRNA ($2^{-\Delta Ct}$) ± SEM, PC1 (n=3-9); PC2 (n=2-7); PC3 (n=3-4). (H) Heatmap representation of biomark analysis of PC1, PC2, PC3 organoids (days 7, 25, 50, 90) for mesoderm lineage (in red); endoderm lineage (blue), trophoblast (yellow) stem cells (black); proliferation (purple); and ectoderm lineage (green).

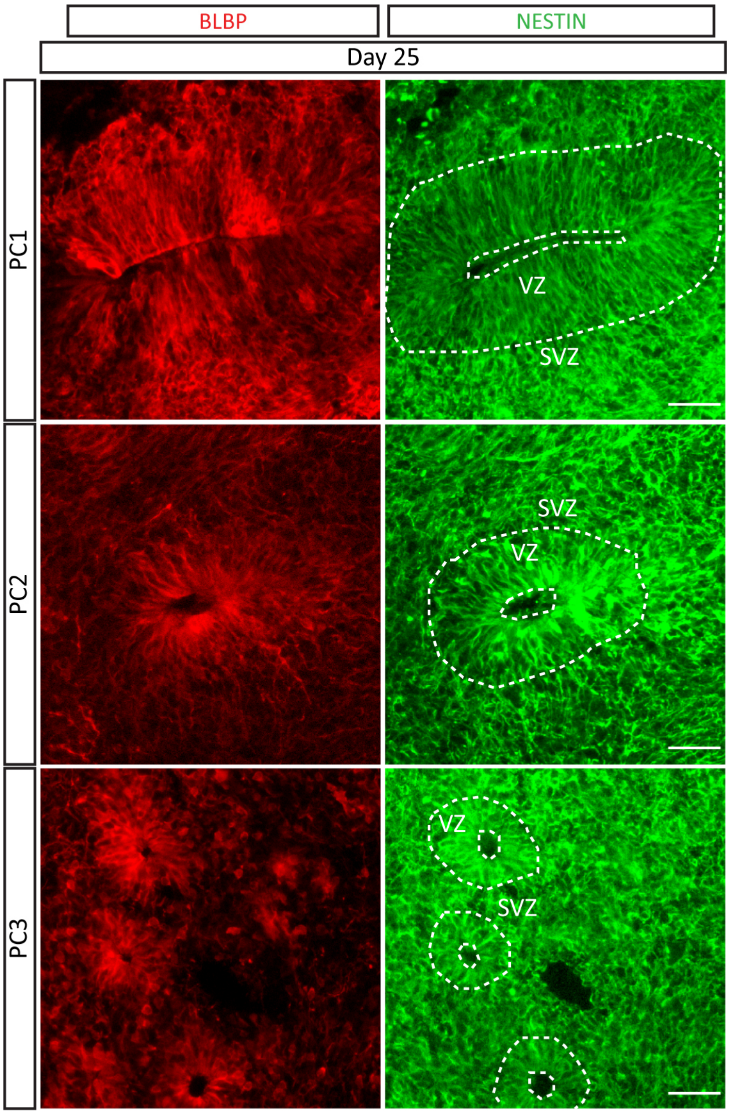


**Figure S2: Cortical organoids at early stages show radial glia, mitotic and neuronal progenitor markers**

Immunofluorescence images showing the expression of the radial glia marker Blbp (red) and the neuronal progenitor marker Nestin (green) at day 25. Scale bars represent 50 μm.

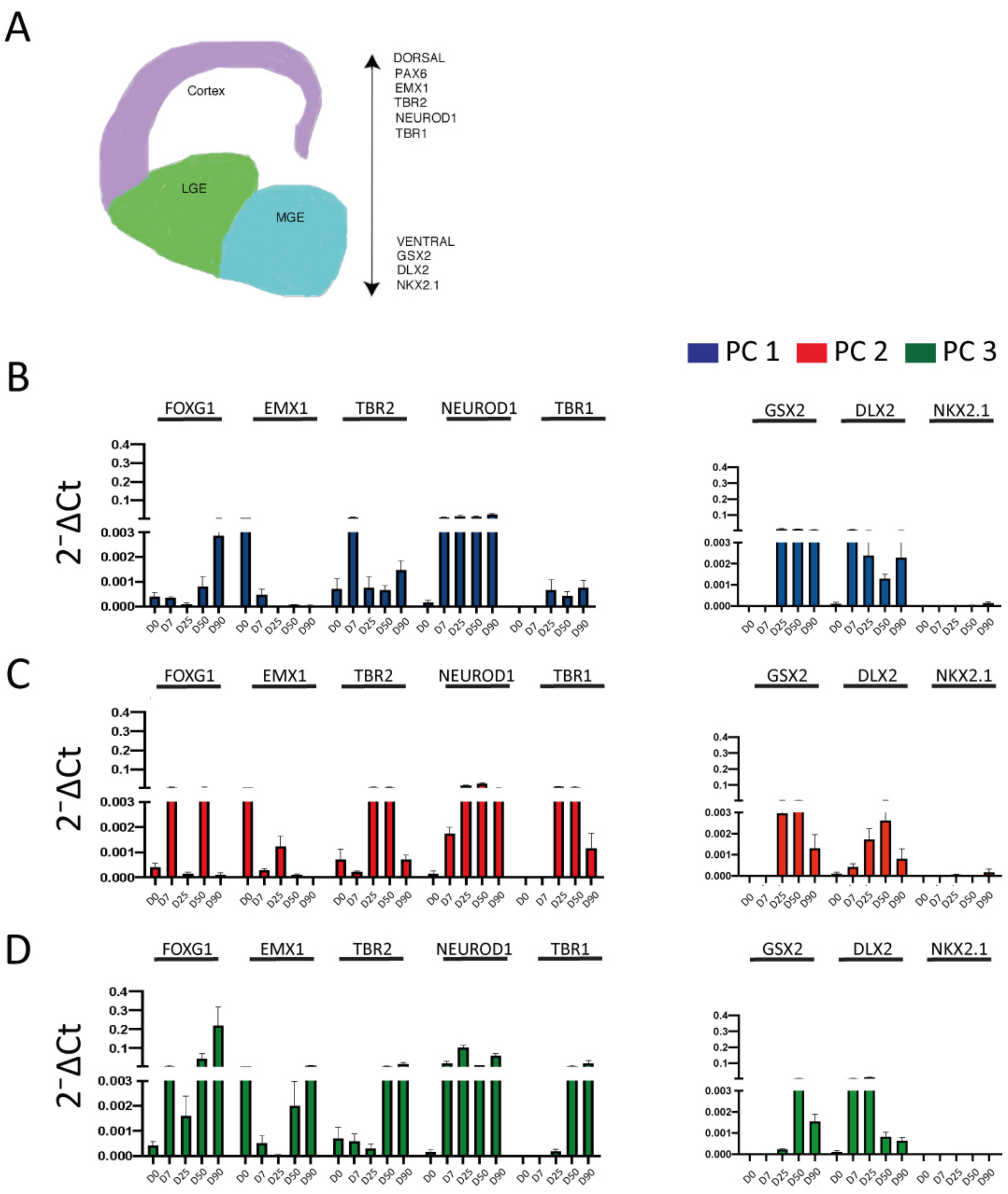


**Figure S3: Dorsal/Ventral pattern of expression of PC1,PC2,PC3-derived organoids**

(A) Cartoon depicting the Dorsal-Ventral axis of the developing telencephalon. (B-D) Biomark analysis of PC1, PC2, PC3 organoids (days 7, 25, 50, 90), for the expression of dorsal: PAX6, FOXG1, EMX1, TBR2, NEUROD1, TBR1; and ventral: GSX2, DLX2, NKX2.1, ISL1, GAD1, GAD2 forebrain-specific genes. Data are shown as absolute normalized amount of mRNA ($2^{-DCt}$) ± SEM, PC1 (n=3-9); PC2 (n=2-7); PC3 (n=3-4) (see Tables S1-S2).

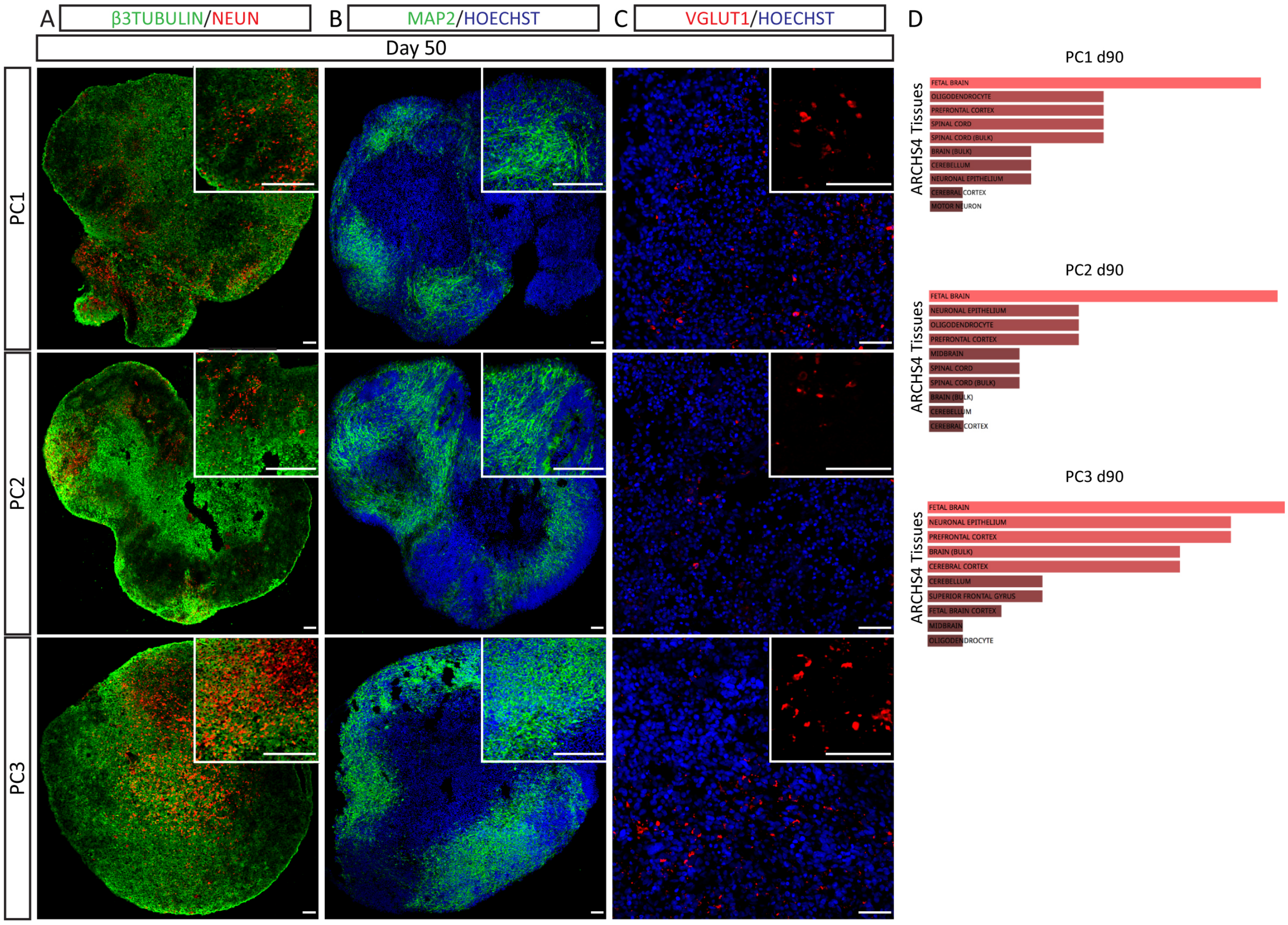


**Figure S4: Late stage organoids express neuronal maturation and glial markers**

(A-C) Immunofluorescence images showing the expression of the neuronal markers β3 tubulin (green; A); NeuN (red; A); Map2 (green; B); VGlut1 (red; C) in day 50 organoids from PC1, PC2 and PC3. Counterstaining of nuclei was done with Hoechst (blue). Scale bars represent 100 μm, and top right insets 400 μm. (D) Top 30 Biomark genes at day 90 for each protocol PC1,PC2, PC3 were analysed for Cell type Analysis: ARCHS4 Tissues (Enrichr website). PC1 d90 : Fetal brain; oligodendrocyte; prefrontal cortex; spinal cord; spinal cord (bulk); brain (bulk); cerebellum; neuronal epithelium; cerebral cortex; motor neuron. PC2 d90: Fetal brain; neuronal epithelium; oligodendrocyte; prefrontal cortex; midbrain; spinal cord; spinal cord (bulk); brain (bulk); cerebellum; cerebral cortex. PC3 d90: Fetal brain; neuronal epithelium; prefrontal cortex; brain (bulk); cerebral cortex; cerebellum; superior frontal gyrus; fetal brain cortex; midbrain; oligodendrocyte.

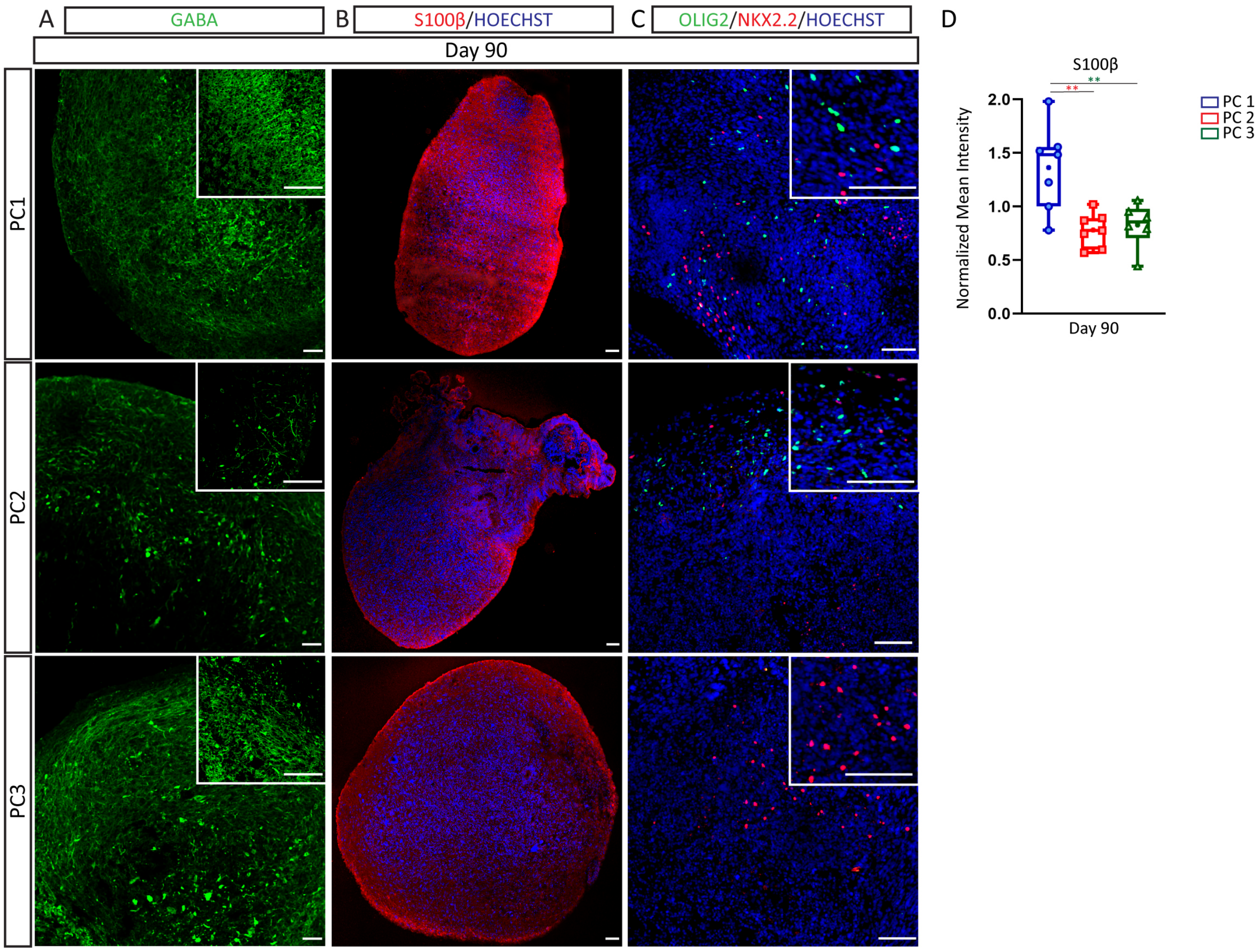


**Figure S5: Wide range of neuronal and glial identities following default conditions of differentiation**

(A-C) Immunofluorescence images showing the expression of the GABAergic neuronal marker GABA (green; A), the astrocytic marker S100β (red; B), and the oligodendrocyte markers Olig2 (green; C); and Nkx2.2 (red; C) in day 90 organoids following PC1, PC2 and PC3 protocols. Counterstaining of nuclei was done with Hoechst (blue). Scale bars represent 50 µm (A), 100 µm (B-C), top right insets 100 µm (B) and 400 µm (C). (C) Box and whiskers graph showing the normalized mean intensity of S100β (C) in PC1, PC2, PC3 organoids at day 90. n=8 sections from 4 organoids per protocol. One way-ANOVA with Tukey post-tests. $^*p < 0.05$, $^{**}p < 0.01$, $^{***}p < 0.001$, $^{****}p < 0.0001$.

**Supplemental Tables**

| | H9 hESC | PC1 | PC2 | PC3 | 9PCW | 20PCW |
|---|---|---|---|---|---|---|
| | | | | | 2 samples | 2 samples |
| **Day 0** | 2 samples | | | | | |
| **Day 7** | | 3 samples | 2 samples | 4 samples | | |
| **Day 25** | | 5 samples | 6 samples | 3 samples | | |
| **Day 50** | | 7 samples | 7 samples | 4 samples | | |
| **Day 90** | | 9 samples | 3 samples | 4 samples | | |

**Table S1: List of organoid samples used for Biomark assay (Fluidigm)**
List of organoid samples, the embryonic stem cell samples H9-WA09, and the primary human embryonic cortical samples used for Biomark analysis. Each sample is a pool of 3-4 independent organoids, from at least 2 independent experiments. PC1 (default differentiation); PC2 (directed differentiation with Wnt activation); PC3 (directed differentiation without Wnt activation).

| | | | |
|---|---|---|---|
| **Reference genes** | GADPH UBC PPIA YWHAZ GUSB 18S | **Ventral Forebrain genes** | GSX2 DLX2 DLX1 DLX5 NKX2.1 |
| **Stem cell genes** | POU5F1 NANOG | **Cortical Neuronal genes** | NEUROD1 BRN2 DAB1 SATB2 TBR1 CTIP2 FEZF2 CUX1 |
| **Proliferative genes and progenitor genes** | MKI67 MCM2 PROM1 | **Gabaergic genes** | GAD1 GAD2 SLC32A1 CALB2 CALB1 STT |
| **Neuronal genes** | NES DCX TUBB3 | **Astrocyte genes** | GFAP S100β SLC1A3 ALDH1L1 GJB6 AQP4 FABP7 SLC1A2 |
| **Meso/Endo/ Trophoderm** | CDX2 SOX17 TBXT KRT14 MIXL1 GSC | **Microglia genes** | Iba1 TREM2 Runx1 TLR4 CD68 |
| **Rostral SNC identity genes** | ZBTB18 ASCL1 DACH1 ARX EPHA7 OTX1 EMX2 LHX2 SIX3 DLX5 ISL1 TH CHAT | **Oligodendrocyte genes** | OLIG2 MBP CNP MAG MOG |
| **Caudal SNC identity genes** | IRX3 EN1 EN2 LMX1A LMX1B FOXA1 FOXA2 IRX1 IRX2 GBX2 SLC6A4 HOXA2 HOXA4 HOXB1 HOXB3 | **Neuronal maturation genes** | MAP2 RBFOX3 PSD95 SLC17A7 CamKIIa GRIA1 GRIN2B |
| **Cortical progenitor genes** | PAX6 FOXG1 EMX1 TBR2 | **Other** | HTT |

**Table S2: List of genes analysed by Biomark assay (Fluidigm)**
Table displaying the reference genes and gene categories tested by Biomark assay in this study.

#### Supplemental Experimental Procedures

##### Maintenance of hESC

Human H9 ES cells were obtained from WiCell at early passage with verified normal karyotype and contamination-free. hESC were regularly checked and maintained mycoplasma free. H9 human embryonic stem cells were cultured both in mTeSR™1 (STEMCELL™ Technologies) on hESC-qualified matrigel-coated dishes and on Mouse Embryonic Fibroblasts-mitomycin C-treated from CD1 (MEF-mt) in the following cell culture media: DMEM/F12 (Gibco) containing 20% KnockOut Serum, 100μM non-essential amino acids (Gibco, 1:100), GlutaMax (Gibco, 1:100), 100μM β-mercaptoethanol (Sigma-Aldrich), 100U/mL penicillin and 100 μg/mL streptomycin (Euroclone) and 10ng/ml FGF2 (R&D Systems). EDTA (0.5mM) or Collagenase (1mg/ml) were used for cell passaging one or twice per week.

##### Protocols for cortical organoid formation

Cortical organoids were generated with few modifications from previous published methods (PC1: (Lancaster et al., 2013); PC2: (Qian et al., 2016); PC3: (Pasca et al., 2015)).

PC1: On day 0 of the differentiation process, H9 hES cells from matrigel-coated dishes with low passage <50 were dissociated into single cell suspension using Accutase. 9.000 cells were plated into each well from a V-bottom non-treated 96 well-plate (Corning) in human ES media with basic fibroblast growth factor (20 ng/ml) and 10μM ROCK inhibitor Y-27632 (STEMCELL™ Technologies). The medium was changed every other day for the first 3 days, then transferred to 6 cm low adhesion plates (Corning) in the same medium without ROCK inhibitor. On day 13 organoids were embedded in Matrigel® as shown before (Lancaster et al., 2013; Lancaster & Knoblich, 2014). Organoids were grown in differentiation medium containing a 1:1 mixture of DMEM/F12 (Gibco) and Neurobasal (Gibco) with N2 supplement (Gibco, 1:200), 100μM non-essential amino acids (Gibco, 1:100), B27 supplement w/o RA (Gibco, 1:100), 50μM 2-βmercaptoethanol (Sigma-Aldrich), 5ng/ml Insulin (Sigma Aldrich, 1:4000). From day 35-onwards B27 supplement with RA (Gibco) was used. Medium was changed every two days and an orbital shaker was used from day 16 onwards.

PC2: H9 hES cells with a passage lower than 50 were dissociated from cultures on MEFs by collagenase treatment (Qian et al., 2016). Suspended colonies were subsequently transferred into ultra-low-attachment 100mm plastic plates (Corning) in hESC medium w/o FGF2. For the first 24 h (day 0), the medium was supplemented with 10 μM of the ROCK inhibitor Y-27632 (STEMCELL™ Technologies). Aggregates were pre-patterned with dual SMAD inhibitors: dorsomorphin (Sigma, 2μM) and A83-01 (Sigma, 2μM) from day 0-5. Aggregates were embedded in Matrigel® from day 5-14. At this stage, two factors, GSK-3β inhibitor CHIR99021 and the TGFβ inhibitor SB-431542, were added from day 5-14. At day 14, Matrigel® embedding was removed and floating spheroids were moved to neural differentiation medium (NM) containing DMEM/F12 (Gibco) with 100U/mL penicillin, 100 μg/mL streptomycin, 2 mM L-Glutamine and supplemented by: B27 supplement w/o RA (Gibco, 1:100), N2 supplement (Gibco, 1:100), 100μM non-essential amino acids (Gibco, 1:100), 100μM β-Mercaptoethanol (Sigma-Aldrich), 2.5μg/ml Insulin (Sigma Aldrich). B27 supplement with RA (Gibco, 1:100), 20 ng/ml BDNF (Peprotech), 20 ng/ml GDNF (Peprotech), 1ng/ml TGF-β, and 0.5mM cAMP were added from day 71-onwards. Organoids were kept on an orbital shaker from day 16 onwards.

PC3: H9 hES cells with a passage lower than 50 were dissociated from MEFs by collagenase treatment (Pasca et al., 2015). Suspended colonies were transferred into ultra-low-attachment 100mm plastic plates (Corning) in hESC medium w/o FGF2. For the first 24 h (day 0), the medium was supplemented with 10 μM ROCK inhibitor Y-27632 (STEMCELL™ Technologies). BMP and TGF-β signaling pathways were inhibited with two small molecules: dorsomorphin (Sigma,10μM) and SB-431542 (Tocris, 10μM) from day 0-6. On the sixth day, the floating spheroids were shifted to neural medium (NM) containing Neurobasal (Gibco), B27 supplement w/o RA (Gibco, 1:50), GlutaMax (Gibco, 1:100), 100U/mL penicillin and100 μg/mL streptomycin. The NM was supplemented with 20ng/ml FGF2 (R&D Systems) and 20ng/ml epidermal growth factor (EGF) (R&D Systems) from day 6-25. 20 ng/ml of BDNF (Peprotech) and NT3 (Peprotech) were added from day 25 till day 43. The organoids were kept on an orbital shaker from day 16 onwards.

##### 3D Organoid tissue preparation and immunofluorescence

Organoids were fixed in 4% paraformaldehyde (PFA) for 45 minutes at 4°C followed by washing in PBS 1X three times for 10 minutes. Tissues were allowed to sink in 15% sucrose O/N at 4°C and then transferred to 30% sucrose (Sigma-Aldrich) for 1-2 days at 4°C. Finally, the organoids were embedded in O.C.T. (O.C.T. compound, VWR BDH Chemicals). Embedded tissue was frozen on dry ice and stored at -80°C until sectioning as 12-15 μm sections using a cryostat (Leica). Cryosections were washed in PBS 1X, permeabilized in 0.5% Triton X-100 in PBS and blocked in 10% Goat Serum + 0.1% Triton X-100 in PBS for 1 hr at room temperature. Primary antibody incubations were performed at 4°C overnight and secondary antibody incubations at room temperature for 1-2 hr in 3% Goat Serum + 0.1% Triton X-100 in PBS O/N at 4°C (see table primary and secondary antibodies below). Secondary antibodies used were goat Alexa Fluor 488, 568 and 647-conjugated goat anti-mouse, -rabbit, -rat IgG (Invitrogen 1:500).

When required antigen retrieval was performed by boiling the slides at 90°C for 15 minutes in citrate buffer 10mM pH=6 prior to the permeabilisation step. Nuclei counterstaining was performed with Hoechst.

| **Primary Antibodies** | **Species** | **Company** | **Catalog Number** | **Dilution** |
|---|---|---|---|---|
| Beta III Tubulin | Rabbit | Covance | PRB-435P | 1:1000 |
| Beta III Tubulin | Mouse | Promega | G7121 | 1:1500 |
| Blbp | Rabbit | Millipore | 16060337 | 1:500 |
| Ctip2 | Rat | Abcam | AB18465 | 1:300 |
| GABA | Rabbit | Sigma | A2052 | 1:1000 |
| GAD67 | Mouse | Millipore | MAB5406 | 1:150 |
| GFAP | Rabbit | Dako | Z0334 | 1:1000 |
| Gsx2 | Rabbit | GeneTex | GTX129390 | 1:250 |
| Iba1 | Rabbit | Wako | 019-19741 | 1:100 |
| Ki67 | Mouse | Cell Signaling | 2586S | 1:800 |
| Ki67 | Rabbit | Abcam | AB16667 | 1:500 |
| Map2 | Mouse | BD | 556320 | 1:1000 |
| Nestin | Mouse | Millipore | 15563322 | 1:200 |
| NeuN | Mouse | Chemicon | MAB377 | 1:100 |
| Nkx2.1 (ttf1-H190) | Rabbit | Santa Cruz | SC53136 | 1:1000 |
| Nkx2.2 | Mouse | Hybridoma Bank | 74.5A5 | 1:100 |
| Olig2 | Rabbit | Millipore | AB9610 | 1:500 |
| Pals1 | Rabbit | Proteitech | 17710-1-AP | 1:500 |
| Pax6 | Mouse | DSHB | 042348 | 1:150 |
| pVimentin | Mouse | MBL | D076-35 | 1:750 |
| S100β | Rabbit | Sigma | ABN59 | 1:100 |
| Satb2 | Mouse | Abcam | AB5152 | 1:500 |
| Sox2 | Mouse | Chemicon | MAB4343 | 1:200 |
| Sox2 | Rabbit | Millipore | AB13970 | 1:200 |
| Tbr1 | Rabbit | Abcam | AB31940 | 1:300 |
| VGlut1 | Mouse | Millipore | 16304824 | 1:300 |

**Widefield microscopy and confocal image acquisition**

Images from fixed and cryosectioned organoids were obtained with a widefield inverted Leica DMI6000B microscope and a confocal TSC SP5 (Leica Microsystems) driven by Las-AF software. The confocal was outfitted with 20x, 40x oil and 63x oil objectives lenses. For excitation 405nm, 488nm, 561nm, 638nm laser lines were used. Acquisition parameters (gain, offset, laser intensity, pinhole dimension) were set with control sections and kept for the acquisition of all conditions. For each immunocytochemistry analysis, at least three different sections of three independent organoids for each differentiation experiment were acquired and the choice of the fields was based on the Hoechst nuclear counterstain.

**Quantification of organoid size**

Organoid size was analysed from 25 and 90 days in vitro organoids. Fixed, cryopreserved and ultrasectioned organoids (12-15 μm sections) were imaged following nuclear staining using Hoechst. Middle sections of organoids were chosen for image acquisition using a Leica DMI6000B microscope (Leica Microsystems) with an air 4x objective driven by Las-AF software. Tile images of the organoids (days 25 and 90) were generated through the Las-AF software and used for further analysis. Image J/Fiji software was used to evaluate the area of the organoid pictures and represented as $mm^2$.

Days 25 and 90 PC1 (n=4); PC2 (n=4); PC3 (n=4). Statistical analysis was performed using a two-way ANOVA with Tukey post-tests, ns= non-significant.

**Quantification of the percentage of Ki67+, Sox2+, Pax6+, Tbr1+, Ctip2+, Satb2+, NeuN+, Iba1+, pVimentin+ cells**

The percentage of positive cells among the total number of cells (Hoechst) of an organoid was determined from immunofluorescence images acquired by confocal TSC SP5 microscope using Fiji/ImageJ quantification tools. Percentage of cells (%) was calculated using a default threshold and generating masks to identify and count particles. Results are shown for PC1, PC2, PC3. n=8 sections from 4 organoids per protocol. Data are expressed as mean values in violin plots.

Statistical analysis was done using one or two way-ANOVA with Tukey post-tests.

**Quantification of the Normalized Mean intensity**
Mean Intensity of an organoid was determined from immunofluorescence images acquired by confocal TSC SP5 microscope using Fiji/ImageJ quantification tools. Mean Intensity values of a given marker was subtracted from control background mean intensity. Normalized Mean Intensity was calculated using an internal control as value 1. Results are shown for PC1, PC2, PC3. n=8 sections from 4 organoids per protocol. Data are expressed as mean values in box and whiskers plots. Statistical analysis was done using one or two way-ANOVA with Tukey post-tests.

**Quantification of the number of normalized VGlut1 puncta per area**
The number of VGlut1 puncta per area of an organoid was determined from immunofluorescence images acquired by confocal TSC SP5 microscope using Fiji/ImageJ quantification tools. Number of puncta was calculated using a default threshold and generating masks to identify and count number of puncta. Number of punta per area was calculated by dividing the number of identified number of puncta by the area. Normalized number of punta per area was calculated using an internal control as value 1. Results are shown for PC1, PC2, PC3. n=8 sections from 4 organoids per protocol. Data are expressed as mean values in violin plots. Statistical analysis was done using two way-ANOVA with Tukey post-tests.

**Quantification of Rosette area and VZ Area**
The area of rosettes and VZ regions was determined from immunofluorescence DAPI/Hoechst images acquired by confocal microscopy using Fiji/ImageJ quantification tools. The Area of the VZ region was calculated by subtracting the internal apical area (Pals1) from the total rosette area. Results are shown for PC1, PC2, PC3. n=8 sections from 4 organoids per protocol. Data are expressed as mean values in violin plots. Statistical analysis was done using one way-ANOVA with Tukey post-tests.

**Human embryonic and adult cortical samples**
Post-mortem human fetal cortical specimens from 9pcw were obtained from University of Cambridge, UK. Embryonic and fetal age was extrapolated based on the date of the mother's last menstruation, ultrasound scans of the fetus in utero, crown-rump length (CRL) and visual inspection. Samples were chilled on ice during dissection and immediately frozen in dry ice and stored at −80 °C for later RNA extraction. All procedures were approved by the research ethical committees and research services division of the University of Cambridge and Addenbrooke's Hospital in Cambridge (protocol 96/85, approved by Health Research Authority, Committee East of England—Cambridge Central in 1996 with amendments in November 2017) in accordance with the Human Tissue Act 2006. RNA from post-mortem human brain specimens at 20pcw was available in the laboratory and obtained according to an established protocol (Onorati, 2014). All documents related to the human fetal samples were submitted to the Ethics Committee of the University of Milano, and ethics approval was obtained on 27 March 2013.
Post-mortem adult cortical samples were obtained from the Harvard Brain Tissue Resource Center (HBTRC) (Belmont, Massachusetts, USA). The Ethics Committee of the University of Milano approved the use of human postmortem samples obtained from HBTRC (Ethics Committee 18.12.13, Ethical Approval 74/14).
Tissue was handled in accordance with ethical guidelines and regulations for the research use of human brain tissue set forth by the National Institute of Health (NIH) (http://bioethics.od.nih.gov/humantissue.html) and the World Medical Association Declaration of Helsinki
(http://www.wma.net/en/30publications/10policies/b3/index.html).

**qRTPCR analysis**
RNA extraction was performed by TRIzol™ Reagent (ThermoFisher scientific). Potential contaminating DNA was removed using Recombinant DNase I (rDNase I, Ambion). RNA quality and concentration was assessed using a Nanodrop. First of all, strand cDNA was synthesized from 500ng of rDNase I-treated total RNA using random primers (Bio-Rad) and iScript cDNA Synthesis Kit (Bio-Rad). Primers for qPCR reaction were designed with either PrimerExpress software (Applied Biosystems), from the websites Invitrogen Oligo perfect or primer bank Harvard Medical School. The efficiency of qPCR amplifications was validated by performing standard curves with four different cDNA dilutions from human embryonic brain tissue with a YWHAZ primer set as normalizer. The quantitative PCR reaction mixtures consisted of SsoFast™ EvaGreen® Supermix 2X (Bio-Rad), 250 nM of primers and 12,5 ng cDNA. The qPCR protocol was performed using a CFX96™ Real-Time System (Bio-Rad). Relative expression levels are expressed as $2^{-\Delta\Delta Ct}$ where $\Delta Ct = Ct_{target} - Ct_{normalizer}$ and $\Delta\Delta Ct = \Delta Ct_{target} - \Delta Ct_{reference}$. Normalizer reference gene was YWHAZ. Primer sequences are described in the table below.
Each sample is a pool of 3-4 independent organoids, from at least 2 independent experiments. Samples used PC1, PC2, PC3 n=3 samples; embryonic cortex (PCW9) n=2, and adult cortex (52 and 63 years old) n=2 samples. Statistical analyses were performed by one-way ANOVA with Tukey post-test of all samples.

| Gene | Primer Forward | Primer Reverse |
|---|---|---|
| 4R TAU | TGCAGATAATTAATAAGAAGCTGGA | GTGTTTGATATTATCCTTTGAGC |
| YWHAZ | ACTTTTGGTACATTGTGGCTTCAA | CCGCCAGGACAAACCAGTAT |

**Biomark HD assay (Fluidigm)**

For high-content qPCR experiments, cDNA was pre-amplified using a 0.2X pool of primers prepared from the same gene expression assays as were used for the Biomark analysis. Pre-amplification allows for multiplexed sequence specific amplification of 96 targets genes. 1.25 µl aliquot of cDNA was pre-amplified in a final volume of 5 µl using 1 µl of PreAmp Master Mix (Fluidigm) and 1.25 µl pooled TaqMan assay mix (0.2X). cDNA went through amplification by denaturing at 95°C for 15 s, and annealing and amplification at 60°C for 4 min for 10 cycles in the Veriti™ 96-Well Thermal Cycler (Thermo Fisher Scientific). After cycling, pre-amplified cDNA was diluted 1:5 by adding 20 µl TE Buffer to the final 5 µl reaction volume for a total volume of 25 µl.

Gene expression experiments were performed using the 96x96 qPCR Dynamic Array microfluidic chips (Fluidigm). A 2.25 µl of amplified cDNA was mixed with 2.5 µl of TaqMan Fast Advanced Master Mix (ThermoFisher scientific) and 0.25 µl of Fluidigm's sample loading agent, then inserted into one of the chip sample inlets. A 2.5 µl aliquot of each 20X TaqMan assay was mixed with 2.5 µl of Fluidigm's assay loading agent and individually inserted into one of the chip assay inlets. Samples and probes were loaded into 96x96 chips using an IFC Controller HX (Fluidigm), then transferred to a BioMark real-time PCR reader (Fluidigm) following manufacturer's instructions. Thermo Fisher Scientific online software was used to design oligonucleotide primers and dye-labelled MGB probes. The list of the 96 TaqMan assays used in this study is provided in the table below and table S2. Samples used in this study are displayed in Table S1. Housekeeping genes used for normalization of the data were YWHAZ and GADPH. Data were analyzed and gene expression was carried out through the use of ΔCT as normalized absolute gene expression analysis: $2^{-\Delta Ct}$ whereby $\Delta Ct = Ct_{target} - Ct_{normalizer}$. Statistical analyses were done using a one-way ANOVA test (Analysis Of Variance), with Tukey post-tests and indicated as * for $p<0.05$; ** for $p<0.01$; *** for $p< 0.001$; **** for $p< 0.0001$.

| Gene Symbol | Gene Name | Cod. | Gene Symbol | Gene Name | Cod. |
|---|---|---|---|---|---|
| GAPDH | glyceraldehyde-3-phosphate dehydrogenase | Hs04420697_g1 | HOXB3 | homeobox B3 | Hs05048382_s1 |
| UBC | ubiquitin C | Hs01871556_s1 | TH | tyrosine hydroxylase | Hs00165941_m1 |
| PPIA | peptidylprolyl isomerase A | Hs04194521_s1 | CHAT | choline O-acetyltransferase | Hs00758143_m1 |
| YWHAZ | tyrosine3-monooxygenase/tryptophan 5-monooxygenase activation protein zeta | Hs01122445_g1 | SLC6A4 | solute carrier family 6 member 4 | Hs00984349_m1 |
| GUSB | glucuronidase beta | Hs00939627_m1 | EMX1 | empty spiracles homeobox 1 | Hs00417957_m1 |
| 18S | Eukaryotic 18S rRNA | Hs99999901_s1 | PAX6 | paired box 6 | Hs00240871_m1 |
| POU5F1 | POU class 5 homeobox 1 pseudogene 5 | Hs01570480_s1 | ZBTB18 | zinc finger and BTB domain containing 18 | Hs00295755_s1 |
| NANOG | Nanog homeobox | Hs02387400_g1 | EOMES | eomesodermin | Hs00172872_m1 |
| MKI67 | marker of proliferation Ki-67 | Hs04260396_g1 | NEUROD1 | neuronal differentiation 1 | Hs01922995_s1 |
| MCM2 | minichromosome maintenance complex component 2 | Hs01091564_m1 | DAB1 | DAB1, reelin adaptor protein | Hs00221518_m1 |
| CDX2 | caudal type homeobox 2 | Hs01078080_m1 | FEZF2 | FEZ family zinc finger 2 | Hs01115572_g1 |
| SOX17 | SRY-box 17 | Hs00751752_s1 | TBR1 | T-box, brain 1 | Hs00232429_m1 |
| TBXT | T brachyury transcription factor | Hs00610080_m1 | BCL11B | B-cell CLL/lymphoma 11B | Hs01102259_m1 |
| KRT14 | keratin 14 | Hs00265033_m1 | Satb2 | SATB homeobox 2 | Hs00392652_m1 |
| MIXL1 | Mix paired-like homeobox | Hs00430824_g1 | Brn2 | POU class 3 homeobox 2 | Hs00271595_s1 |
| GSC | goosecoid homeobox | Hs00418279_m1 | Cux1 | cut like homeobox 1 | Hs00738851_m1 |
| PROM1 | prominin 1 | Hs01009259_m1 | GFAP | glial fibrillary acidic protein | Hs00909233_m1 |
| NES | Nestin | Hs04187831_g1 | S100β | S100 calcium binding protein B | Hs00902901_m1 |
| DCX | Doublecortin | Hs00167057_m1 | SLC1A3 | solute carrier family 1 member 3 | Hs00904823_g1 |
| TUBB3 | tubulin beta 3 class III | Hs00801390_s1 | ALDH1L1 | aldehyde dehydrogenase 1 family member L1 | Hs01003842_m1 |
| EMX2 | empty spiracles homeobox 2 | Hs00244574_m1 | FABP7 | fatty acid binding protein 7 | Hs00361424_g1 |
| OTX1 | orthodenticle homeobox 1 | Hs00951099_m1 | SLC1A2 | solute carrier family 1 member 2 | Hs01102423_m1 |
| SIX3 | SIX homeobox 3 | Hs00193667_m1 | GJB6 | gap junction protein beta 6 | Hs00922742_s1 |
| LHX2 | LIM homeobox 2 | Hs00180351_m1 | AQP4 | aquaporin 4 | Hs00242342_m1 |
| ARX | aristaless related homeobox | Hs00292465_m1 | MAG | myelin associated glycoprotein | Hs01114387_m1 |
| EPHA7 | EPH receptor A7 | Hs01033006_m1 | MOG | myelin oligodendrocyte glycoprotein | Hs01555268_m1 |
| GSX2 | GS homeobox 2 | Hs00370195_m1 | MBP | myelin basic protein | Hs00921945_m1 |
| ASCL1 | achaete-scute family bHLH transcription factor 1 | Hs04187546_g1 | CNP | 2',3'-cyclic nucleotide 3'phosphodiesterase | Hs00263981_m1 |
| DLX2 | distal-less homeobox 2 | Hs00269993_m1 | OLIG2 | oligodendrocyte lineage transcription factor 2 | Hs00300164_s1 |
| DLX1 | distal-less homeobox 1 | Hs00698288_m1 | MAP2 | microtubule associated protein 2 | Hs00258900_m1 |
| DLX5 | distal-less homeobox 5 | Hs00193291_m1 | RBFOX3 | RNAbinding protein, fox-1 homolog 3 | Hs01370654_m1 |
| FOXG1 | forkhead box G1 | Hs01850784_s1 | PSD95 | discs large MAGUK scaffold protein 4 | Hs01555373_m1 |

| | | | | | |
|---|---|---|---|---|---|
| ISL1 | ISL LIM homeobox 1 | Hs00158126_m1 | SLC17A7 | solute carrier family 17 member 7 | Hs00220404_m1 |
| NKX2-1 | transcription termination factor 1 | Hs00201121_m1 | CamK2a | calcium/calmodulin dependent protein kinase II alpha | Hs00947041_m1 |
| DACH1 | dachshund family transcription factor 1 | Hs00974297_m1 | GRIA1 | glutamate ionotropic receptor AMPA type subunit 1 | Hs00181348_m1 |
| IRX3 | iroquois homeobox 3 | Hs01124217_g1 | GRIN2B | glutamate ionotropic receptor NMDA type subunit 2B | Hs01002012_m1 |
| EN1 | engrailed homeobox 1 | Hs00154977_m1 | Iba1 | allograft inflammatory factor 1 | Hs00610419_g1 |
| EN2 | engrailed homeobox 2 | Hs00171321_m1 | Runx1 | runt related transcription factor 1 | Hs02558380_s1 |
| LMX1A | LIM homeobox transcription factor 1 alpha | Hs00898455_m1 | CD68 | CD68 molecule | Hs02836816_g1 |
| LMX1B | LIM homeobox transcription factor 1 beta | Hs00158750_m1 | TREM2 | triggering receptor expressed on myeloid cells 2 | Hs00219132_m1 |
| FOXA1 | forkhead box A1 | Hs04187555_m1 | TLR4 | toll like receptor 4 | Hs00152939_m1 |
| FOXA2 | forkhead box A2 | Hs05036278_s1 | CALB2 | calbindin 2 | Hs00242372_m1 |
| IRX1 | iroquois homeobox 1 | Hs00411782_m1 | SLC32A1 | solute carrier family 32 member 1 | Hs00369773_m1 |
| IRX2 | iroquois homeobox 2 | Hs01383002_m1 | GAD2 | glutamate decarboxylase 2 | Hs00609534_m1 |
| GBX2 | gastrulation brain homeobox 2 | Hs00230965_m1 | GAD1 | glutamate decarboxylase 1 | Hs01065893_m1 |
| HOXA2 | homeobox A2 | Hs00534579_m1 | CALB1 | calbindin 1 | Hs01077197_m1 |
| HOXA4 | homeobox A4 | Hs01573270_m1 | STT | somatostatin | Hs00356144_m1 |
| HOXB1 | homeobox B1 | Hs00157973_m1 | HTT | huntingtin | Hs00918174_m1 |

**Jaccard similarity**
Jaccard similarity index was computed between the lists of the top 100 differentially expressed genes in the CTX compared to LGE and MGE at the indicated time (ranked by the sum of the adjusted p-value of pairwise comparisons between CTX and LGE or MGE, from Bocchi et al. 2021) and the lists of the top 30 highly expressed genes in the Biomark assay in the different protocols at the indicated time of differentiation (mean of 3 replicates).

**PCA, heatmaps and back-to-back analysis**
All statistical analyses and graphical representations were conducted using R v.3.4.2 using core packages as *cluster*, *factoextra, pcr, tibble* and *ggplot2*.
Gene clustering was performed through hierarchical approach; co-profiled samples were clustered together by using the euclidean method, while for co-expressed genes Pearson Correlation method was performed. In both cases, complete linkage method was used, as from suggestions by Fluidigm official Faqs (https://www.fluidigm.com/faq/ge-54).
Top three Principal Components from gene expression matrix were selected to carry out top 10 up/downregulated genes.

**Brain Atlas Database**
Heatmap representation of the top 10 up biomark genes from PC1,PC2 and PC3 organoids at day 90 was done from human brain samples (amigdala, basal ganglia, cerebellum, cerebral cortex, hippocampus, hypothalamus, midbrain, olfactory region, pons and medulla, thalamus) data obtained from Brain Atlas database: https://www.proteinatlas.org/humanproteome/brain.

**Venn diagram and Cell Type representation from biomark Analysis**
The representation of the Venn diagram and the Cell Type Analysis (ARCHS4) representation of the top 30 biomark genes of PC1,PC2,PC3 at day 90 was performed using the enrichr website.

**Statistical analysis**

The data obtained from different analyses carried out in this work were processed with GraphPad Prism v.7 software. Statistical analysis was done following a one-way ANOVA or two-way ANOVA test (Analysis Of Variance), with Tukey post-tests and indicated as * for $p<0.05$; ** for $p<0.01$; *** for $p< 0.001$; **** for $p< 0.0001$.
